# Arv1 interacts with and regulates the first step of GPI biosynthesis in *Candida albicans*

**DOI:** 10.1101/2024.11.16.623920

**Authors:** Monika Bharati, Harshita Saini, Neha Thakran, Yatin Kumar, Usha Yadav, Shailja Shefali, Sunyna Saun, Aaisha Anzar, Sneha Sudha Komath

## Abstract

The ubiquitous *ARV1* gene shows significant functional conservation across eukaryotes. In humans, it is implicated in early onset epileptic encephalopathy. Evidence suggests that the phenotypes manifested in affected patients are probably due to the deficiency in expression of cell surface GPI anchored proteins. *S. cerevisiae* Arv1 is proposed to be the elusive GPI flippase that delivers the GPI intermediate from the cytoplasmic face to the luminal side of the ER for further elaboration by the first mannosyltransferase of the pathway. Human and fungal *ARV1* complement *S. cerevisiae ARV1*. Overexpressing some of the GPI-*N*-acetylglucosamine transferase (GPI-GnT) subunits rescues the null strain of *S. cerevisiae ARV1*. In mammals and in *T. brucei* Arv1 co-immunoprecipitates with one or more subunits of the GPI-GnT. Based on these reports we hypothesized a cross-talk for *ARV1* with the GPI biosynthetic pathway in the human pathogenic fungus, *C. albicans*. Using super resolution radial fluctuation (SRRF) analysis for co-localization, co-immunoprecipitation assays, and acceptor-photobleaching Förster resonance energy transfer (FRET) studies, we show that *C. albicans* Arv1 physically interacts with the GPI-GnT. It also transcriptionally regulates the expression of the GPI-GnT genes to control the GPI biosynthetic pathway via its very first step. Overexpression of one of the GPI-GnT subunits, CaGpi19, in *C. albicans ARV1* null strain rescues its cold-sensitive growth, azole sensitivity, cell wall phenotype and GPI-GnT activity. Thus, our results suggest extensive interactions between Arv1 and GPI biosynthesis in *C. albicans*.

## Introduction

Arv1 is a lipid transporter required for sterol homeostasis. It was first identified in *S. cerevisiae* as a gene that could restore viability in a strain lacking *ARE1/2* (acyl-coenzyme A:cholesterol O-acyltransferase (<u>A</u>CAT) <u>r</u>elated <u>e</u>nzyme 1/2) only when *ARE2* was reintroduced [1]. *S. cerevisiae*

*arv1* mutants exhibit a large number of phenotypes: slow growth at permissible temperatures (30 °C), cold shock as well as heat shock sensitivity, altered sterol distribution along with defects in Erg11 levels and accumulation of lanosterol (substrate of Erg11), sensitivity to nystatin and azole drugs, and defects in phospholipid and sphingolipid biosynthesis [1–4]. Mammalian, plant, and fungal (*C. albicans* as well as *C. glabrata*) *ARV1* homologs complement a *S. cerevisiae arv1Δ*, suggesting a significant degree of functional conservation across eukaryotes [1,5–7]. In *C. albicans*, as in *S. cerevisiae*, Arv1 has a physical interaction with Erg11, a crucial enzyme in the ergosterol biosynthetic pathway and the target of azole drugs [8].

Arv1 is also implicated in glycosylphosphatidylinositol (GPI) biosynthesis in several eukaryotes. In humans, splice variants and point mutants of *ARV1* reduce surface level GPI-anchored protein (GPI-AP) expression and produce disease phenotypes that appear to overlap with those observed in patients with inherited GPI deficiencies [9–11]. Arv1 has been shown to physically interact with the first step of GPI biosynthesis in mammals as well as in *T. brucei* [12,13]. The *S. cerevisiae arv1Δ* strain is rescued either partially or completely by the expression of *GPI1, GPI2, GPI3,* and *ERI1*, all encoding different subunits of the GPI-*N*-acetylglucosaminyltransferase (GPI-GnT), an ER membrane-localized enzyme that catalyzes transfer of *N*-acetylglucosamine (GlcNAc) from UDP-GlcNAc to phosphatidylinositol (PI) and initiates the process of GPI biosynthesis [2]. The effect of overexpressing genes coding for the remaining two subunits of the GPI-GnT, *GPI15* and *GPI19*, have not been studied.

The GlcNAc-PI formed on the cytoplasmic face of the ER membrane at the end of this step is sequentially de-*N*-acetylated and then acylated in two different enzyme-catalyzed reactions to produce GlcN(acyl)-PI. The *S. cerevisiae arv1Δ* strain mentioned above accumulates this latter GPI intermediate, suggesting that GPI biosynthesis in this strain is stalled at the end of the third step [2]. In the normal course, in wild type cells, the acylated GPI intermediate would have been flipped into the ER lumen, after which a series of enzyme-dependent mannosylation and ethanolamine phosphate (EtNP) transfer reactions would occur, giving rise to Manα1-6(EtNP)Manα1-2(EtNP)Manα1-6(EtNP)Manα1-4GlcN-(acyl)inositol-phospholipid, the complete GPI precursor (for a detailed review on GPI biosynthesis in yeast see, [14]). It would then be covalently attached by a GPI transamidase (GPIT) to the C-terminal end of various proteins that have the appropriate GPI attachment signal sequence to produce an array of GPI-anchored proteins (GPI-APs). The GPI-APs would have their lipid tails remodeled and their carbohydrate core further modified as they are transported out of the ER, and via the Golgi, to their final destinations in the outer leaflet of the plasma membrane (PM) and/or the cell wall for their function. Thus, as expected, the *S. cerevisiae arv1Δ* strain mentioned above is deficient in the surface expression of GPI-APs as well.

Taken together, the above information brings us to the obvious question, and the main focus of this manuscript: does Arv1 interact with the GPI biosynthetic pathway and with the GPI-GnT in particular in the pathogenic human fungus, *C. albicans*? Previous results from our lab indicate that *C. albicans GPI19*, encoding one of the six GPI-GnT subunits, is transcriptionally co-activated with *ERG11* [15]. Downregulating either of the genes brings down expression levels of the other and activation of either one activates the other. The sensitivity to azole drugs is directly correlated to levels of Erg11 in the *C. albicans* GPI-GnT mutants [15,16]. Given that CaArv1 physically interacts with Erg11, as mentioned above, it is possible to hypothesize that it might also interact with or affect the *C. albicans* GPI-GnT.

Hence, we investigated the effect of deleting *CaARV1* on the GPI-GnT and the first step of GPI biosynthesis in *C. albicans*. We show that a homologous null strain of *CaARV1* has reduced GPI-GnT activity and, therefore, reduced levels of GPI-APs at the cell surface. These cells also show altered cell wall biogenesis and sensitivity to ketoconazole. Overexpression of *CaGPI19* in these cells reverses these phenotypes. Interestingly, CaArv1 co-localizes to the ER along with CaGpi2, Cagpi19 and CaGpi3, all subunits of the GPI-GnT, suggesting that CaArv1 could possibly interact with the GPI-GnT in *C. albicans*. Acceptor-photobleaching FRET too supports such a hypothesis.

## RESULTS

The strains and plasmids used in this study are listed in Table 1 A&B.

**Table 1:**
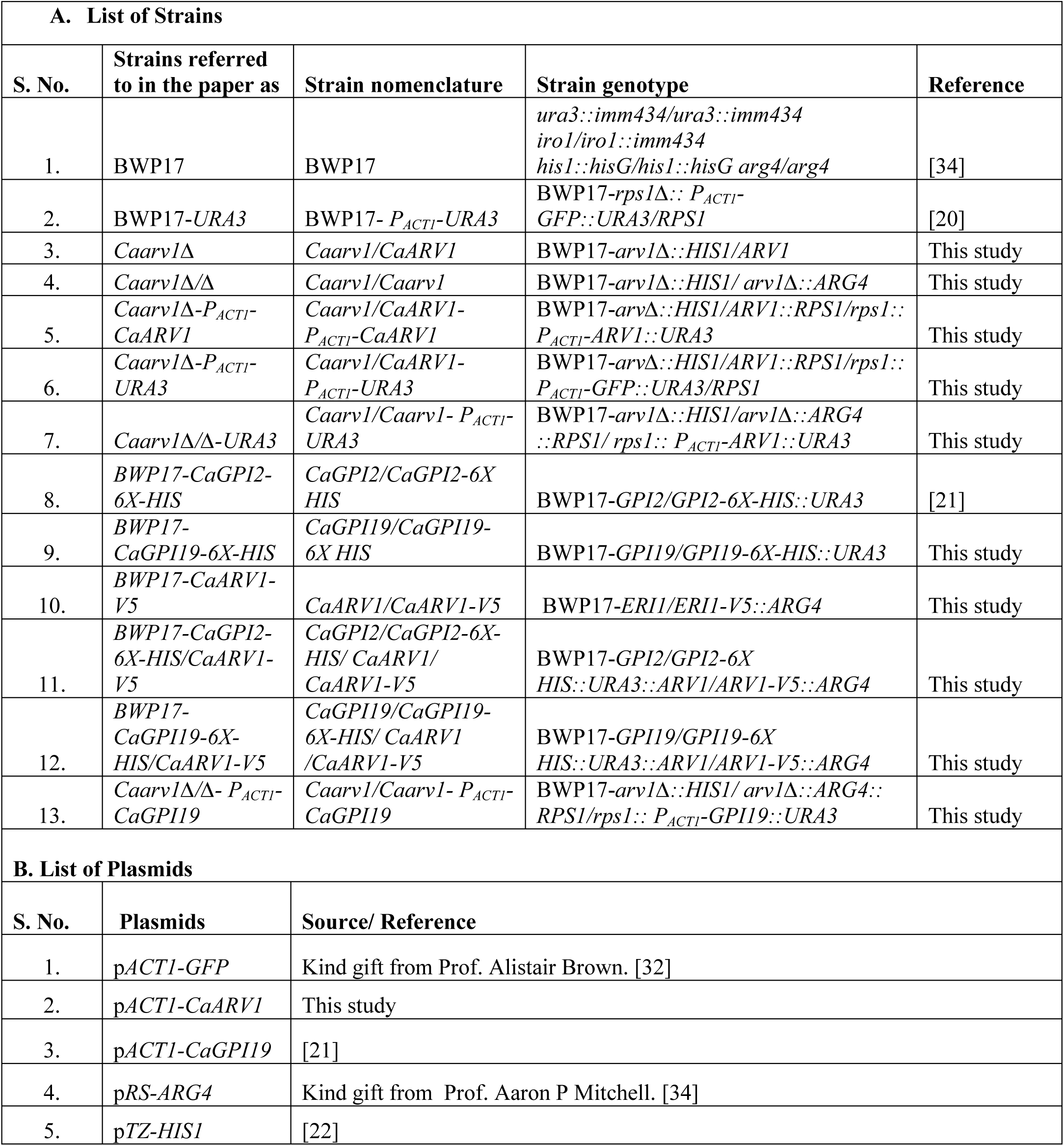
*C. albicans* strains and the plasmids used in the study.

| **A. List of Strains** |  |  |  |  |
| --- | --- | --- | --- | --- |
| **S. No.** | **Strains referred to in the paper as** | **Strain nomenclature** | **Strain genotype** | **Reference** |
| 1. | BWP17 | BWP17 | *ura3::imm434/ura3::imm434*<br>*iro1/iro1::imm434*<br>*his1::hisG/his1::hisG arg4/arg4* | [34] |
| 2. | BWP17-URA3 | BWP17- *P<sub>ACT1</sub>*-URA3 | BWP17-*rps1Δ:: P<sub>ACT1</sub>-*<br>*GFP::URA3/RPS1* | [20] |
| 3. | *Caarv1Δ* | *Caarv1/CaARV1* | BWP17-*arv1Δ::HIS1/ARV1* | This study |
| 4. | *Caarv1Δ/Δ* | *Caarv1/Caarv1* | BWP17-*arv1Δ::HIS1/ arv1Δ::ARG4* | This study |
| 5. | *Caarv1Δ-P<sub>ACT1</sub>-*<br>*CaARV1* | *Caarv1/CaARV1-*<br>*P<sub>ACT1</sub>-CaARV1* | BWP17-*arvΔ::HIS1/ARV1::RPS1/rps1::*<br>*P<sub>ACT1</sub>-ARV1::URA3* | This study |
| 6. | *Caarv1Δ-P<sub>ACT1</sub>-*<br>*URA3* | *Caarv1/CaARV1-*<br>*P<sub>ACT1</sub>-URA3* | BWP17-*arvΔ::HIS1/ARV1::RPS1/rps1::*<br>*P<sub>ACT1</sub>-GFP::URA3/RPS1* | This study |
| 7. | *Caarv1Δ/Δ-URA3* | *Caarv1/Caarv1- P<sub>ACT1</sub>-*<br>*URA3* | BWP17-*arv1Δ::HIS1/arv1Δ::ARG4*<br>*::RPS1/ rps1:: P<sub>ACT1</sub>-ARV1::URA3* | This study |
| 8. | *BWP17-CaGPI2-*<br>*6X-HIS* | *CaGPI2/CaGPI2-6X*<br>*HIS* | BWP17-*GPI2/GPI2-6X-HIS::URA3* | [21] |
| 9. | *BWP17-*<br>*CaGPI19-6X-HIS* | *CaGPI19/CaGPI19-*<br>*6X HIS* | BWP17-*GPI19/GPI19-6X-HIS::URA3* | This study |
| 10. | *BWP17-CaARV1-*<br>*V5* | *CaARV1/CaARV1-V5* | BWP17-*ERI1/ERI1-V5::ARG4* | This study |
| 11. | *BWP17-CaGPI2-*<br>*6X-HIS/CaARV1-*<br>*V5* | *CaGPI2/CaGPI2-6X-*<br>*HIS/ CaARV1/*<br>*CaARV1-V5* | BWP17-*GPI2/GPI2-6X*<br>*HIS::URA3::ARV1/ARV1-V5::ARG4* | This study |
| 12. | *BWP17-*<br>*CaGPI19-6X-*<br>*HIS/CaARV1-V5* | *CaGPI19/CaGPI19-*<br>*6X-HIS/ CaARV1*<br>*/CaARV1-V5* | BWP17-*GPI19/GPI19-6X*<br>*HIS::URA3::ARV1/ARV1-V5::ARG4* | This study |
| 13. | *Caarv1Δ/Δ- P<sub>ACT1</sub>-*<br>*CaGPI19* | *Caarv1/Caarv1- P<sub>ACT1</sub>-*<br>*CaGPI19* | BWP17-*arv1Δ::HIS1/ arv1Δ::ARG4::*<br>*RPS1/rps1:: P<sub>ACT1</sub>-GPI19::URA3* | This study |
| **B. List of Plasmids** |  |  |  |  |
| **S. No.** | **Plasmids** | **Source/ Reference** |  |  |
| 1. | p*ACT1-GFP* | Kind gift from Prof. Alistair Brown. [32] |  |  |
| 2. | p*ACT1-CaARV1* | This study |  |  |
| 3. | p*ACT1-CaGPI19* | [21] |  |  |
| 4. | p*RS-ARG4* | Kind gift from Prof. Aaron P Mitchell. [34] |  |  |
| 5. | p*TZ-HIS1* | [22] |  |  |

### Heterozygous and homozygous null strains of *ARV1* in *C. albicans* grow normally, and are sensitive to ketoconazole

The heterozygous strain of *C. albicans ARV1* (*Caarv1/CaARV1*, referred to as *Caarv1Δ* from here on) was generated in BWP17 strain background by disrupting one allele using a *HIS1* marker, and the homozygous null (*Caarv1/Caarv1*, referred to as *Caarv1Δ/Δ*) was generated by disrupting the second allele using an *ARG4* marker. A reintegrant strain was generated by introducing a single copy of the gene into the heterozygous *Caarv1Δ* strain at the *RPS1* locus using the constitutively active promoter, P*_ACT1_* (*Caarv1/CaARV1/*P*_ACT1_-CaARV1*; referred to as *Caarv1Δ-*P*_ACT1_-CaARV1* from here on). The strains were confirmed using PCR (Figure S1 A (i)-(iii)). Because the reintegrant strain was generated using *URA3*, a gene that exhibits positional effects [17], one copy of it was introduced at the same locus in the other two strains to generate, *Caarv1Δ-*P*_ACT1_-URA3,* and *Caarv1Δ/Δ-*P*_ACT1_-URA3* and confirmed by PCR (Figure S1 A (iv)-(v)). BWP17-P*_ACT1_-URA3* was used as the wild type control [18]. These strains are referred to as *Caarv1Δ-URA3*, *Caarv1Δ/Δ-URA3*, and BWP17*-URA3,* respectively, from here on.

As can be seen from Figure 1A, *CaARV1* transcript levels are significantly reduced in *Caarv1Δ-URA3* while it is restored to wild type levels in *Caarv1Δ-*P*_ACT1_-CaARV1*. No significant amounts of *CaARV1* transcripts are detected in *Caarv1Δ/Δ-URA3*. None of the strains show any growth defects in either solid (Figure S1B) or liquid media at 30 °C (Figure 1B(i); Table 2). Both *Caarv1Δ-URA3,* and *Caarv1Δ/Δ-URA3* grow slowly in the presence of the azole drug, ketoconazole (0.15 μg/ml), as compared to the wild type control (Figure 1B(ii); Table 2). This is consistent with what is expected of *C. albicans* strains deficient in *ARV1* [8]. It is also consistent with the downregulation of mRNA transcripts of *CaERG11* that we observe in *Caarv1Δ-URA3,* and *Caarv1Δ/Δ-URA3* (Figure 1C(i)) and the small but significant drop in ergosterol levels in *Caarv1Δ/Δ*-*URA3* (Figure 1C(ii)). *Caarv1Δ/Δ-URA3* also exhibits a cold-sensitive growth phenotype (Figure 1D; Table 2). These phenotypes are reversed in the reintegrant strain, *Caarv1Δ-* P*_ACT1_-CaARV1*, proving that they are specific to *CaARV1*.

**Figure 1:**
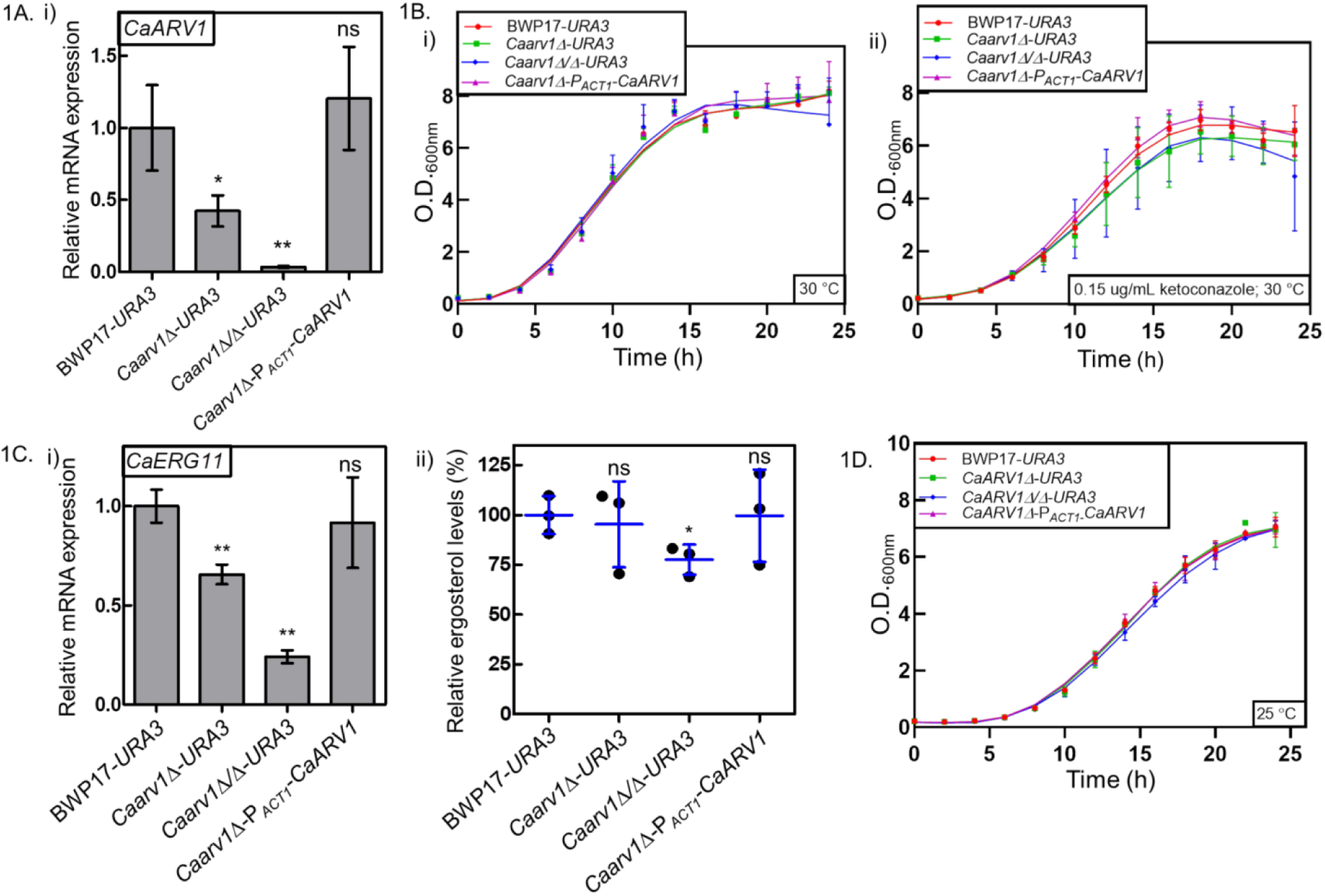
Confirmation, growth and azole sensitivity of mutant strains of *CaARV1*. **A.** (i**) Transcript analysis of *CaARV1* in the mutant strain.** *CaARV1* mRNA transcripts are reduced in *Caarv1Δ-URA3* and *Caarv1Δ/Δ-URA3* in comparison to BWP17-*URA3* and restored in *Caarv1Δ-*P*_ACT1_-CaARV1.* **B. Growth of *CaARV1* mutants in the absence and presence of ketoconazole.** Secondary cultures were grown at 30 °C in SD Ura^-^ liquid media using cells from a primary culture corresponding to O.D._600nm_ ∼ 0.2. Growth was monitored every 2 h, **(i)** in the absence of ketoconazole and **(ii)** in the presence of 0.15 µg/ml ketoconazole. The doubling times obtained are given in Table 2. **C. Sterol biosynthetic pathway is perturbed in *CaARV1* mutant strains. (i) Transcript levels of *CaERG11* reduce in the heterozygous and null strain and are restored in the *CaARV1* reintegrant.** mRNA levels of *CaERG11* in *Caarv1Δ-URA3, Caarv11Δ/Δ-URA3* and *Caarv1Δ-*P*_ACT1_-CaARV1* in comparison to BWP17-*URA3*. *GAPDH* was used as the internal standard. **(ii) Ergosterol levels decrease in *Caarv1Δ/Δ-URA3*.** GC-MS analysis was shown here is average of three independent sets done. **D. *Caarv1Δ/Δ-URA3* is cold sensitive.** Secondary cultures were grown at 25 °C in SD Ura^-^ liquid media using cells from a primary culture corresponding to O.D._600nm_ ∼ 0.2. Growth was monitored after every 2 h. The doubling times obtained are given in Table 2. The statistical significance was calculated using the unpaired T-test with Welch’s correction. ‘*’ represents P-values calculated relative to the wild type strain; ‘ns’ implies non-significant. All experiments were repeated thrice with independent cultures.

**Table 2:**
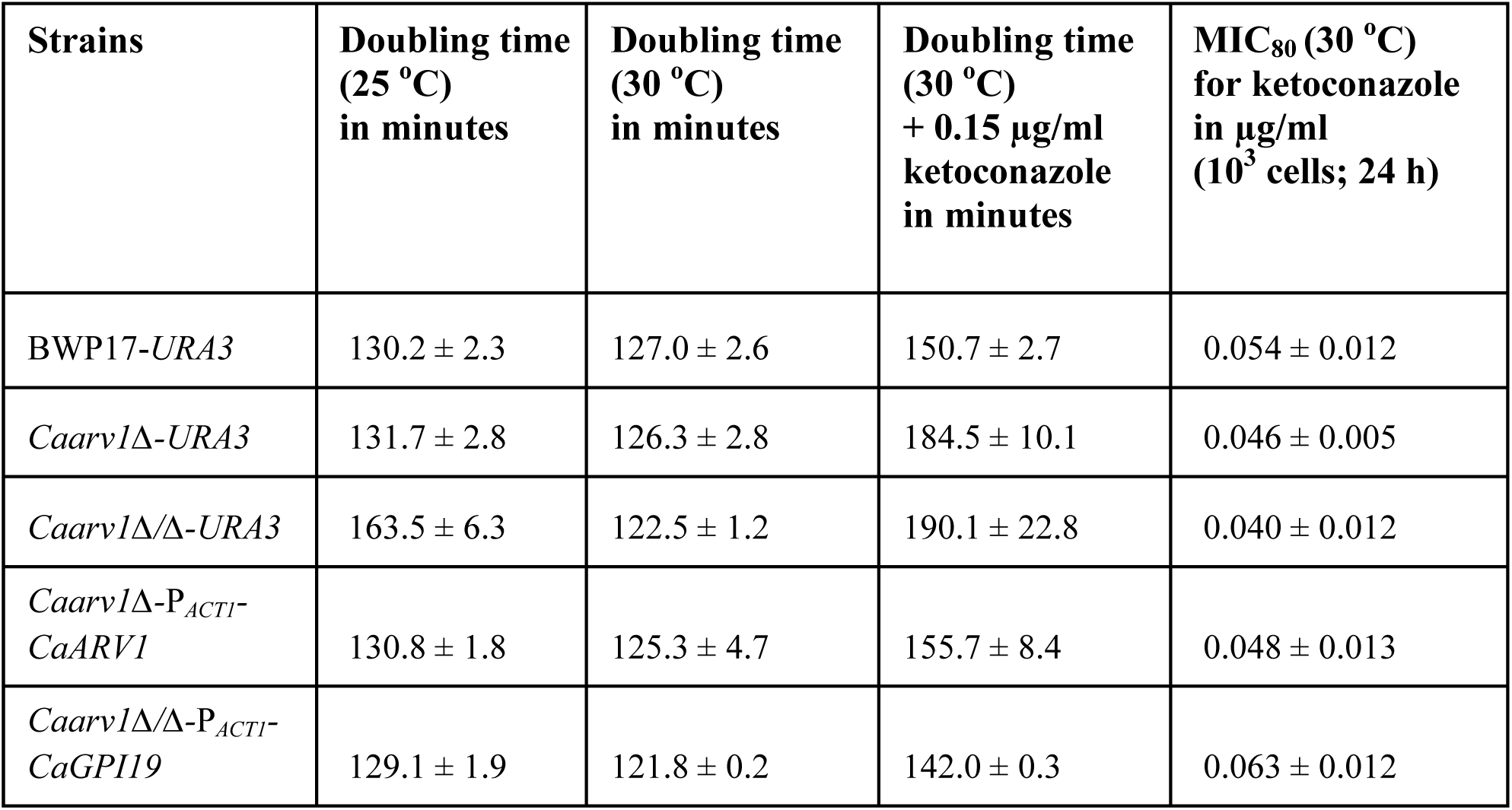
Doubling times of the *C. albicans* strains used in this study in the absence and presence of ketoconazole and the MIC_80_ values of ketoconazole for them.

### Heterozygous and homozygous null strains of *CaARV1* have thicker cell walls and are defective in filamentation via the cAMP-PKA pathway

*S. cerevisiae arv1Δ* is sensitive to calcofluor white (CFW), a cell wall perturbing agent, and accumulates chitin [2]. Hence, the *CaARV1* deficient strains were tested for sensitivity to CFW. We see no obvious sensitivity to CFW for either *Caarv1Δ-URA3* or *Caarv1Δ/Δ-URA3* (Figure 2A) although we do observe significant accumulation of chitin in the cell wall (Figure 2B (i)-(ii)). *Caarv1Δ-URA3* and *Caarv1Δ/Δ-URA3* are also significantly hypofilamentous at 37 °C in Spider medium (Figure 2C (i)-(iii)) as well as in DMEM + 5% CO_2_ as compared to BWP17-*URA3* (Figure 2D (i)-(iii)). Since these media activate filamentation in *C. albicans* via the cAMP-PKA pathway, it would suggest that this pathway is specifically downregulated in these strains [19]. The cell wall and filamentation phenotypes are reversed in *Caarv1Δ-*P*_ACT1_-CaARV1*, proving that these phenotypes too are *CaARV1*-specific.

**Figure 2:**
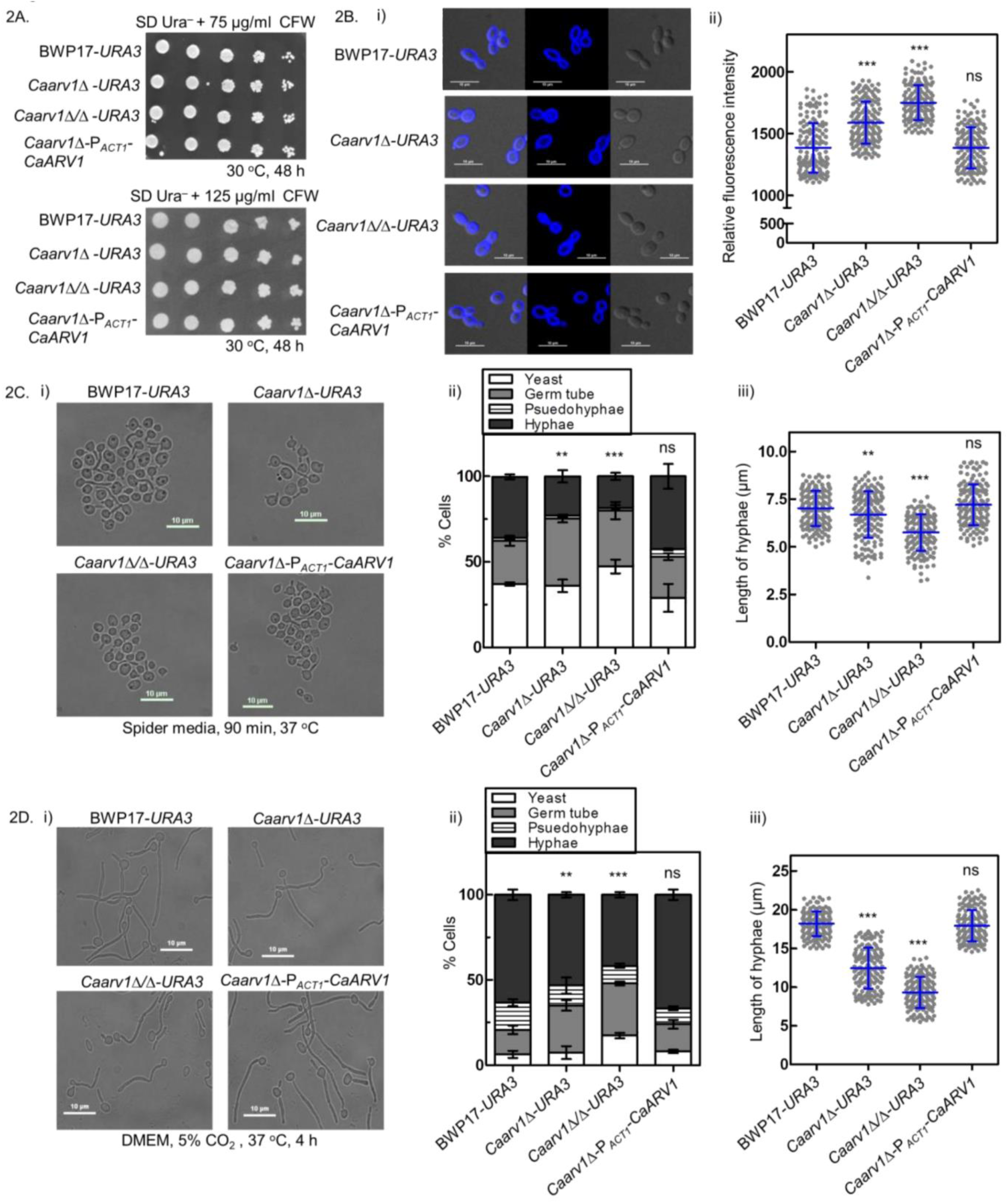
Heterozygous and homozygous null strains of *CaARV1* have thicker cell walls and are defective in filamentation: **A. *CaARV1* deletion mutants are not sensitive to CFW**. Mid-log phase cells were taken at O.D._600nm_ ∼ 0.2 and five-fold serially diluted for spotting on SD Ura^-^-agar containing 75 and 125 µg/ml CFW followed by incubation at 30 °C. **B. *CaARV1* deficient strains accumulate chitin in the cell wall.** Cells grown in SD Ura^-^ broth were stained with 100 µg/ml CFW. **(i)** Representative images are shown. **(ii)** Fluorescence intensities were quantified from 50 cells of each set. Averages and standard deviations are from three independent sets of experiments. The statistical significance was calculated using the unpaired T-test with Welch’s correction. **C. *CaARV1* deficient cells are hypofilamentous via the Ras-dependent pathway.** Primary cultures of the above mentioned strains grown in SD Ura^-^ broth were inoculated in Spider media and incubated for 90 min at 37 °C for hyphal induction. **(i)** Representative images for each strain are shown here. **(ii)** Numbers of yeast cells, germ tube, psuedohyphae and hyphae were counted manually for 100 cells. Three independent sets of experiments were done. Statistical analysis shown here is for hyphal cells only and was calculated using two-way ANOVA with Tukey’s multiple comparison test. **(iii)** Hyphae length were quantified for 50 cells in each set and shown in these plots along with their averages and standard deviations. The statistical significance was calculated using the unpaired T-test with Welch’s correction. **D. *CaARV1* deletion strains are hypofilamentous via the CO_2_-dependent pathway.** Primary cultures of the above mentioned strains grown in SD Ura^-^ broth were used to inoculate DMEM medium at 37 °C in the presence of 5% CO_2_ for 4 h. **(i)** Representative images for the different strains shown. **(ii)** Numbers of yeast cells, germ tube, pseudohyphae and hyphae were counted manually for 100 cells. Three independent sets of experiments were done. Statistical analysis shown here is for hyphal cells only and was calculated using two-way ANOVA with Tukey’s multiple comparison test. **(iii)** Hyphae length were quantified for 50 cells in each set and shown in these plots along with their averages and standard deviations. The statistical significance was calculated using the unpaired T-test with Welch’s correction. P-values calculated for the mutant strains relative to the wild type strain are represented as ‘*’.

### *CaARV1* deletion strains are defective in GPI biosynthesis

GPI biosynthesis is downregulated in both *Caarv1Δ*-*URA3* and *Caarv1Δ/Δ*-*URA3* and restored to wild type levels in *Caarv1Δ-*P*_ACT1_-CaARV1*. This is true whether we monitor GPI-GnT activity (Figure 3A) or estimate the cell surface levels of Als5, a GPI-AP of *C. albicans* (Figure 3B (i)-(ii)). Since the *Caarv1Δ/Δ*-*URA3* strain shows reduced ergosterol levels, as shown above, we wondered whether ergosterol deficiency in the membrane is responsible for the reduced GPI-GnT activity. To test this, the PMF fraction from *Caarv1Δ/Δ*-*URA3* was supplemented with different amounts of ergosterol in the GPI-GnT assay, but this causes no significant alteration in the enzyme activity (Figure S1C).

**Figure 3:**
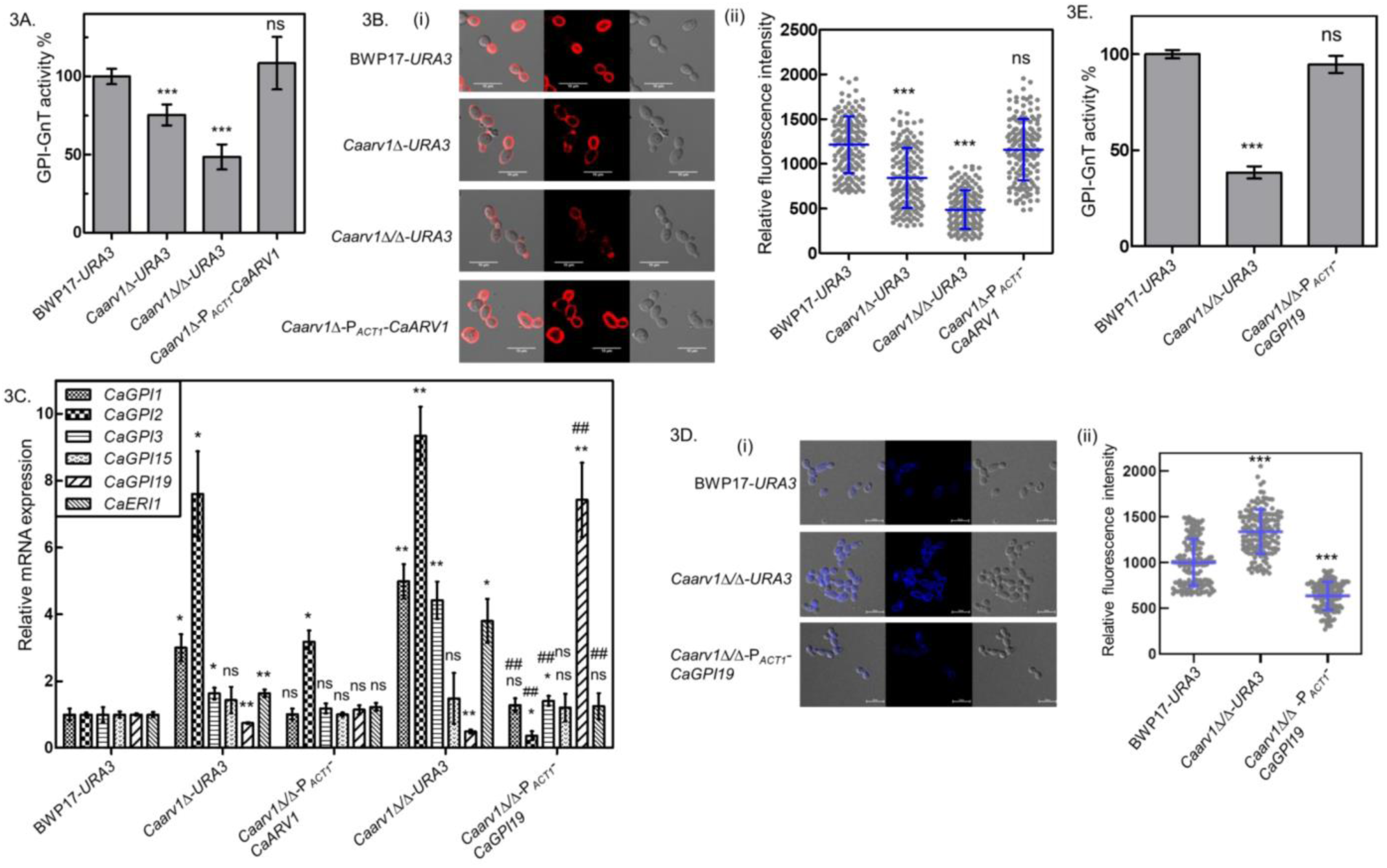
*CaARV1* deletion strains are defective in GPI biosynthesis: **A. GPI-GnT activity is downregulated in *CaARV1* mutants.** PMF from late log phase cells grown in SD Ura^-^ broth were incubated with UDP-[6-^3^H]GlcNAc. The glycolipids extracted were resolved on HPTLC plates and detected/ quantified using Bioscan AR2000. The data are shown relative to BWP17-*URA3*. Averages and standard deviation are from three independent sets. **B. The levels of GPI-APs on the cell surface decrease in *CaARV1* deletion strains.** Mid-log phase cells grown in SD Ura^-^ broth were immunostained with anti-Als5 antibody (GPI-AP). **(i)** Representative images of the mentioned strains are shown here. **(ii)** Relative fluorescence intensity of 50 cells each from 3 independent experiments are shown. Averages and standard deviations are calculated from three independent sets. **C. Transcript levels of GPI-GnT subunits in *CaARV1* mutants. (i)** Transcript levels analysis of expression of different GPI-GnT subunits in the different strains used in this study are shown. **D. *CaGPI19* overexpression reverses the cell wall defects of *Caarv1Δ/Δ-URA3.*** CFW staining was used to detect chitin accumulation in the strains. **(i)** Representative images of the strains are shown. **(ii)** Mid-log phase cells grown in SD Ura^-^ broth were stained with 100 µg/ml CFW. Fluorescence intensities from 50 cells each from 3 independently performed experiments and shown along with averages and standard deviations. **E. GPI-GnT activity is restored in *Caarv1Δ/Δ-* P*_ACT1_-CaGPI19***. PMF generated from cells grown in SD Ura^-^ broth were incubated with UDP-[6-^3^H]GlcNAc as donor substrate for the GPI-GnT activity. The statistical significance was calculated using the unpaired T-test with Welch’s correction. The P-values calculated for the *CaARV1* mutant strains relative to the wild type strain are represented using * while those with reference to *Caarv1Δ/Δ-URA3* are shown using *^#^*.

Next, we examined whether inter-subunit transcriptional regulations within the *C. albicans* GPI-GnT are perturbed by the loss of *CaARV1*, which too would affect GPI-GnT activity [16,18,20,21]. As can be seen in Figure 3C, there is a clear downregulation of *CaGPI19* in the *Caarv1Δ/Δ-URA3* which is reversed in *Caarv1Δ-*P*_ACT1_-CaARV1*. The levels of *CaGPI15* are unaltered while those of the other genes are upregulated. It is interesting to note that the mutually negative regulations between *CaGPI19* and *CaGPI2*, that we previously reported [20], is maintained in the *CaARV1* deficient strains as well. Further, the transcriptional cross-talk within the GPI-GnT is specific to the loss of *CaARV1* and is restored in the cells of *Caarv1Δ*-P*_ACT1_-CaARV1*.

### *CaGPI19* overexpression rescues the *Caarv1Δ/Δ*-*URA3* strain

Not only are the transcript levels of *CaGPI19* reduced in *Caarv1Δ/Δ*-*URA3*, as explained above, many of its phenotypes are similar to the phenotypes exhibited by *CaGPI19* deficient strains. The latter too have reduced *ERG11* levels, are azole sensitive, and accumulate higher amounts of chitin in the cell wall [15,20,22]. Hence, *CaGPI19* was overexpressed in the *Caarv1Δ/Δ* strain using the P*_ACT1_*promoter (Figure S2A). This strain, *Caarv1Δ/Δ*-P*_ACT1_-CaGPI19*, shows reversal of azole sensitivity (Figure S2B; Table 2) and a restoration in its cell wall phenotype relative to *Caarv1Δ/Δ*-*URA3* (Figure 3D (i)-(ii)). In addition, it reverses the cold-sensitivity of the parent strain (Figure S2C; Table 2) and restores GPI-GnT activity (Figure 3E) along with the transcript levels of the other GPI-GnT subunits (Figure 3C), suggesting that a genetic interaction exists between *CaARV1* and *CaGPI19*.

The filamentation phenotype of *CaARV1* deficient mutants is distinct from that of the *CaGPI19* mutants. Heterozygous and conditional null strains of *CaGPI19* are hyperfilamentous and overexpression of *CaGPI19* in a heterozygous strain of *CaGPI19* reverses its hyperfilamentation in Spider medium [15,21]. We also know that this phenotype is controlled by the increased expression levels of *CaGPI2* in these strains, which respond to the inter-subunit cross-talk within the GPI-GnT [21], as described in the previous section. *Caarv1Δ/Δ*-*URA3* also shows upregulation of *CaGPI2* (Figure 3C) and yet is hypofilamentous in Spider medium (Figure 2C). This suggests that the filamentation defect is not directly related to the altered expression levels of the GPI-GnT subunits, but is a result of the loss of *CaARV1* specifically. Not surprisingly, therefore, overexpression of *CaGPI19* in *Caarv1Δ/Δ*-*URA3* does not reverse the filamentation phenotype; instead, it further exacerbates it (Figure S2D-E).

### CaArv1 co-localizes and physically interacts with subunits of the GPI-*N*-acetylglucosamine transferase (GPI-GnT)

In order to determine whether a physical interaction occurs between CaArv1 and the GPI-GnT, we needed to first confirm that CaArv1 localizes to the ER and with the GPI-GnT specifically. For this, CaArv1 was tagged with V5 at its C-terminus in cells of the wild type strain, BWP17 (Figure S3A(i)). The tagging does not alter the GPI-GnT activity of the strains (Figure S3 B), suggesting that it does not alter their function.

Co-localization studies were carried out using super resolution radial fluctuation (SRRF) analysis. CaArv1 significantly co-localizes with the ER tracker dye in these experiments (Figure 4A(i)). It also co-localizes with three GPI-GnT subunits, CaGpi2, CaGpi3, and CaGpi19 at specific regions within the ER (Figure 4A(ii)-(iv)). Pull-down assays show that CaArv1-V5 comes down onto Ni-NTA beads when either CaGpi2, or CaGpi19 is tagged with His_6_ at the C-terminus (Figure 4B (i)-(ii)). As can be seen from this figure, the converse is also true. When CaArv1-V5 is immunoprecipitated onto Protein A-Sepharose beads with anti-V5 antibody, both CaGpi2-His_6_ and CaGpi19-His_6_ are co-precipitated with it(Figure 4B(iii)-(iv)). The C-terminal His_6_-tag does not affect the function of CaGpi19 (Figure S3B) nor that of CaGpi2 [21]. CaArv1-V5 can also be co-immunoprecipitated with CaGpi3 when anti-CaGpi3 antibodies are used (Figure 4B(v)).

**Figure 4:**
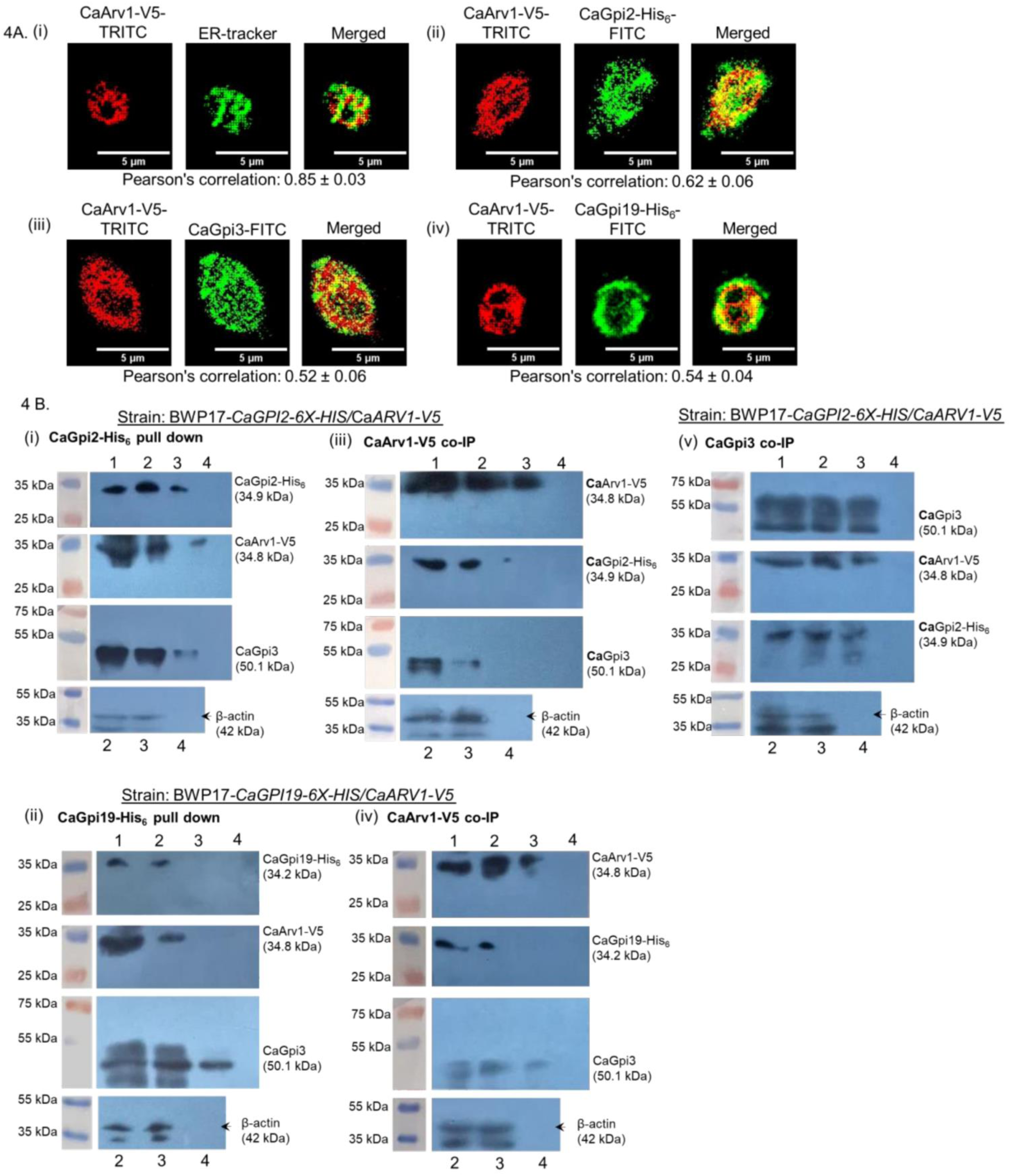
ER localization of CaArv1 and its interaction with GPI-GnT subunits. **A. CaArv1 resides at ER membrane and co-localizes with GPI-GnT subunits.** (i)-(iv) Representative images for each pair used for the co-localization studies are shown here. V5-tagged CaArv1 was immunostained with anti-V5 antibody, ER membrane was stained with ER tracker, CaGpi2-His_6_ and CaGpi19-His_6_ were immunostained with anti-His_6_ antibody, and CaGpi3 was detected with anti-CaGpi3 antibody. SRRF analysis was performed to improve the resolution. The Pearson’s correlation coefficient was calculated using the Nikon NIS elements software; Data of 5 cells each from three independent sets were taken for calculating averages with standard deviations. **B. CaArv1 comes down with CaGpi2, CaGpi19 and CaGpi3 in pull down assays.** (i)-(ii) are representative western blots from experiments where CaGpi2-His_6_ and CaGpi19-His_6_, respectively, were pulled down on Ni-NTA beads, (iii)-(iv) are representative western blots from experiments where CaArv1-V5 was immunoprecipitated with anti-V5 antibodies on Protein A-Sepharose beads, and (v) represents western blots from experiments where CaGpi3 was immunoprecipitated with anti-CaGpi3 antibodies. To detect the proteins, CaArv1-V5 was immunostained with anti-V5 antibody, CaGpi2-His_6_ and CaGpi19-His_6_ were immunostained with anti-His_6_ antibody and CaGpi3 was detected with anti-CaGpi3 antibody. Given their overlapping sizes (their expected molecular sizes are shown in the figure in parenthesis), the samples were run on different SDS-PAGE gels for each pull down assay and western blotting performed with the antibodies mentioned in the figure. β-actin was used as the loading control. In each gel, Lanes-1: elute from beads; 2: Input pre-IP/pull-down; 3: Input post-IP/pull-down; 4: Beads control (Ni-NTA/ protein A-Sepharose beads washed, then eluted with SDS-PAGE sample buffer).

Acceptor photobleaching-fluorescence resonance energy transfer (AP-FRET) experiments also suggest that CaArv1 is located in close proximity to at least three GPI-GnT subunits, CaGpi2, CaGpi19 and CaGpi3 (Figure 5A-C). The calculated distances from the FRET data suggest that CaArv1 is located closer to CaGpi2 (spatial distance, *r* = 5.9 ± 0.9 *nm*) and CaGpi19 (*r* = 5.7 ± 0.8 *nm*) than to CaGpi3 (*r* = 6.3 ± 0.7 *nm*). We previously showed that CaRas1 physically interacts with CaGpi2 and activates the GPI-GnT [18,21]. Hence, this was used as the positive control (*r* = 5.3 ± 0.6 *nm*) (Figure 5D). Further, to determine the specificity of the interaction, we examined the interaction of CaArv1 with CaGpi8, a subunit of the GPI-transamidase required for the attachment of complete GPI precursors to proteins. No energy transfer is observed between the two proteins (*r* > 10 *nm*) (Figure 5E). Thus, it appears that a specific physical interaction exists between CaArv1 and one or more subunits of the GPI-GnT.

**Figure 5:**
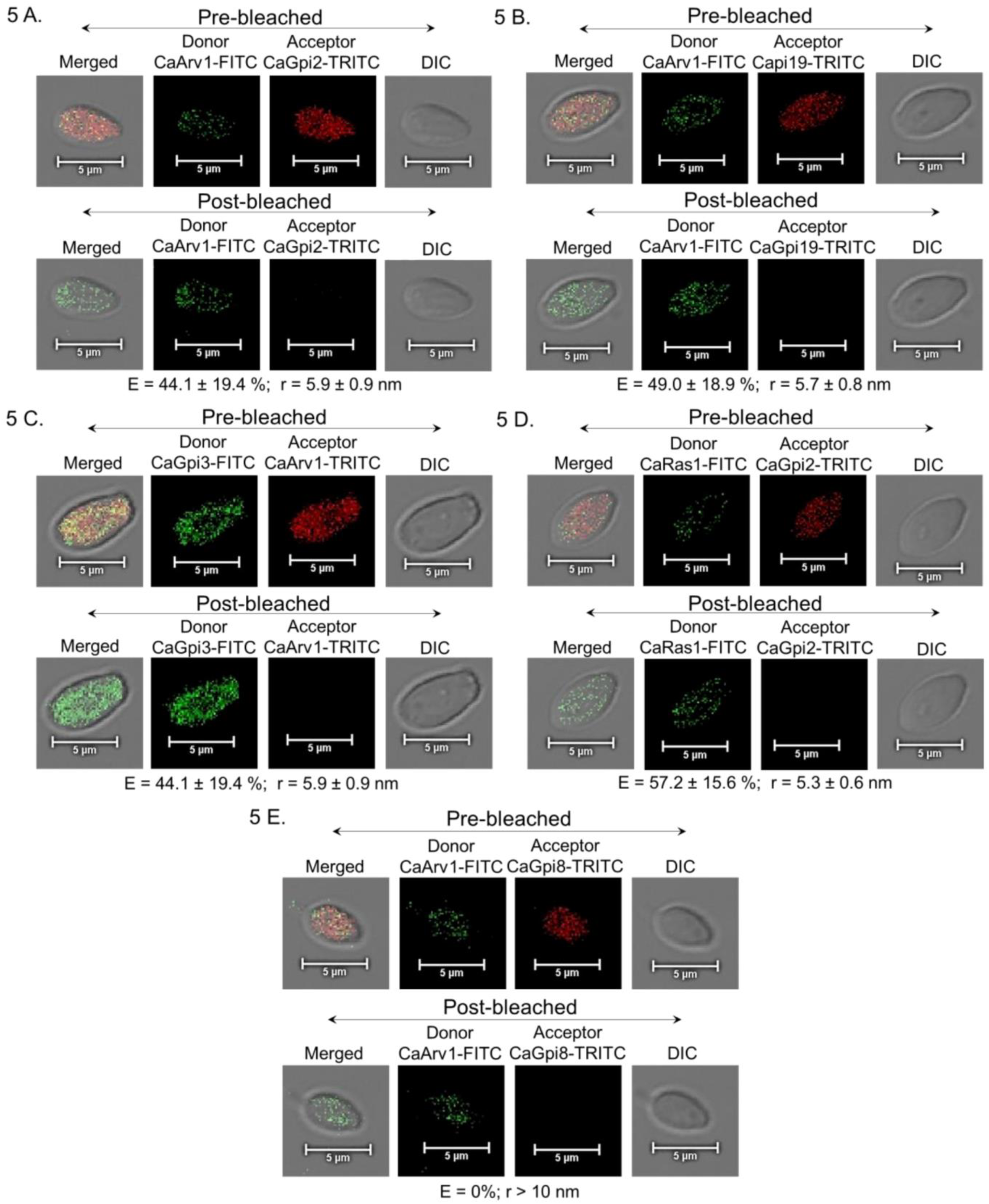
CaArv1 physically interacts with GPI-GnT subunits. CaGpi2-His_6_ and CaGpi19-His_6_ and CaArv1-V5 were probed using appropriate primary antibodies (anti-His_6_ or anti-V5). CaGpi3 and CaGpi8 were detected using custom-synthesized polyclonal antibodies. These were then probed with goat anti-mouse IgG-FITC and goat anti-rabbit IgG-TRITC to generate fluorescence donor-acceptor pairs for AP-FRET experiments as explained in the methodology section. Representative images for each FRET pair with fluorescence intensities pre- and post-bleaching are shown for the donor-acceptor pairs: **A.** CaArv1(FITC)-CaGpi2(TRITC); **B.** CaArv1(FITC)-CaGpi19(TRITC); **C.** CaGpi3(FITC)-CaArv1(TRITC); **D.** CaRas1(FITC)-CaGpi2(TRITC), positive control; and **E.** CaArv1(FITC)-CaGpi8(TRITC), negative control. FRET efficiency and the estimated donor-acceptor distance are presented below each panel. Four ROIs from 4 cells in each set were taken for the calculations. The experiment was repeated thrice using independently cultured cells.

## DISCUSSION

The results presented here confirm that the heterozygous and homozygous null strains of *CaARV1* generated in the BWP17 strain background of *C. albicans* are sensitive to ketoconazole. Similar sensitivity of *CaARV1* null mutants to other azole drugs have been reported in literature [5,8,23]. This is also in line with the downregulation of *CaERG11* and the reduction in levels of ergosterol that we observed, especially in the homozygous null strain. However, whether the extent of downregulation of CaErg11, the target of azoles, is alone sufficient for the sensitivity to azoles is debatable.

*S. cerevisiae arv1Δ* cells are also sensitive to azole drugs, a phenotype which can be reversed by the expression of *CaARV1* in this strain [4,5]. They accumulate a different set of sterol intermediates and possess only slightly lesser than normal levels of ergosterol [4,23,24]. Altered sterol homeostasis as well as alterations in the organization and structure of the PM and ER, are reported when *ARV1* is deleted in either *S. cerevisiae* or *C. albicans* [4,5,24]. It also alters the pleiotropic drug response (PDR) pathway in *arv1Δ* cells which can be reversed by overexpression of CaArv1 [5,8]. Similarly, it is well known that the localization and function of at least some of the multi-drug resistance (MDR) efflux pumps of *C. albicans* are ergosterol-dependent [25–27].

Our results also show that the homozygous null strain of *CaARV1* was hypofilamentous in both Spider medium and in DMEM + 5% CO_2_. While the former medium activates the cAMP-PKA pathway in a Ras-dependent manner, the latter activates it in a Ras-independent manner [19]. Hence, it appears that the cAMP-PKA signaling, controlling a major hyphal morphogenetic pathway in *C. albicans*, is reduced due to the loss of CaArv1. Suppression of cAMP-PKA signaling too has been associated with sensitivity to azoles in *C. albicans* [28]. Further, the results presented here show a clear alteration in the *C. albicans* cell wall in a *CaARV1* dependent manner. Increased deposition of chitin in the cell wall of *C. albicans* strains is often a response to cell wall damage and alterations in the cell wall integrity which are known to affect drug susceptibilities [29]. Thus, it may be argued that a combination of factors could be dictating the response of *Caarv1Δ/Δ*-*URA3* cells to azole drugs.

Inter-subunit transcriptional cross-talk at the first step of the GPI biosynthetic pathway is unique to *C. albicans* and has not so far been reported to exist in any other organism [19]. The fact that the transcript levels are restored to near normal in the *CaARV1* reintegrant strain along with the GPI-GnT activity, is suggestive of the fact that *CaARV1* is specifically involved in this process. Mammalian ARV1 was recently reported to be involved in upregulation of an early step of GPI biosynthesis (resulting in accumulation of GlcN-PI, at the very least) along with upregulation of specific precursor GPI-anchored proteins, although no transcriptional alteration of GPI biosynthetic genes was observed [12]. While the exact nature of how CaArv1 may be involved in regulating the GPI-GnT genes in *C. albicans* requires further investigation, we did observe alterations in the transcript levels of all GPI-GnT genes other than *CaGPI15.* Specifically, *CaGPI19* transcripts are significantly downregulated in the homozygous null strain of *CaARV1*. We know that *CaGPI19* is co-regulated with *ERG11* [15,20], hence, it seemed natural to ask whether overexpression of *CaGPI19* in the *CaARV1* null strain would restore *ERG11* levels as well as the other phenotypes of the strain. Interestingly, it not only reversed *ERG11* levels and the ketoconazole sensitivity of the cells but also their cell wall phenotype as well as the GPI-GnT activity. The filamentation phenotype of the *CaARV1* null strain does not appear to directly arise from alterations in the GPI-GnT activity, since restoration of GPI-GnT activity did not reverse the filamentation. It also appears to be independent of the expression levels of the GPI-GnT subunits, since *CaGPI2* is overexpressed in the *CaARV1* null strain and yet produces hypofilamentation, contrary to what would have been expected if the GPI-GnT subunit was involved [21]. What is also interesting to note is that the mutually negative transcriptional regulation between *CaGPI2* and *CaGPI19* [20] persists in the *CaARV1* null cells. Indeed, the fact that *CaGPI19* overexpression further suppresses the hypofilamentous phenotype of the *CaARV1* null strain further confirms this fact. These results are summed up in a model shown in Figure 6.

**Figure 6:**
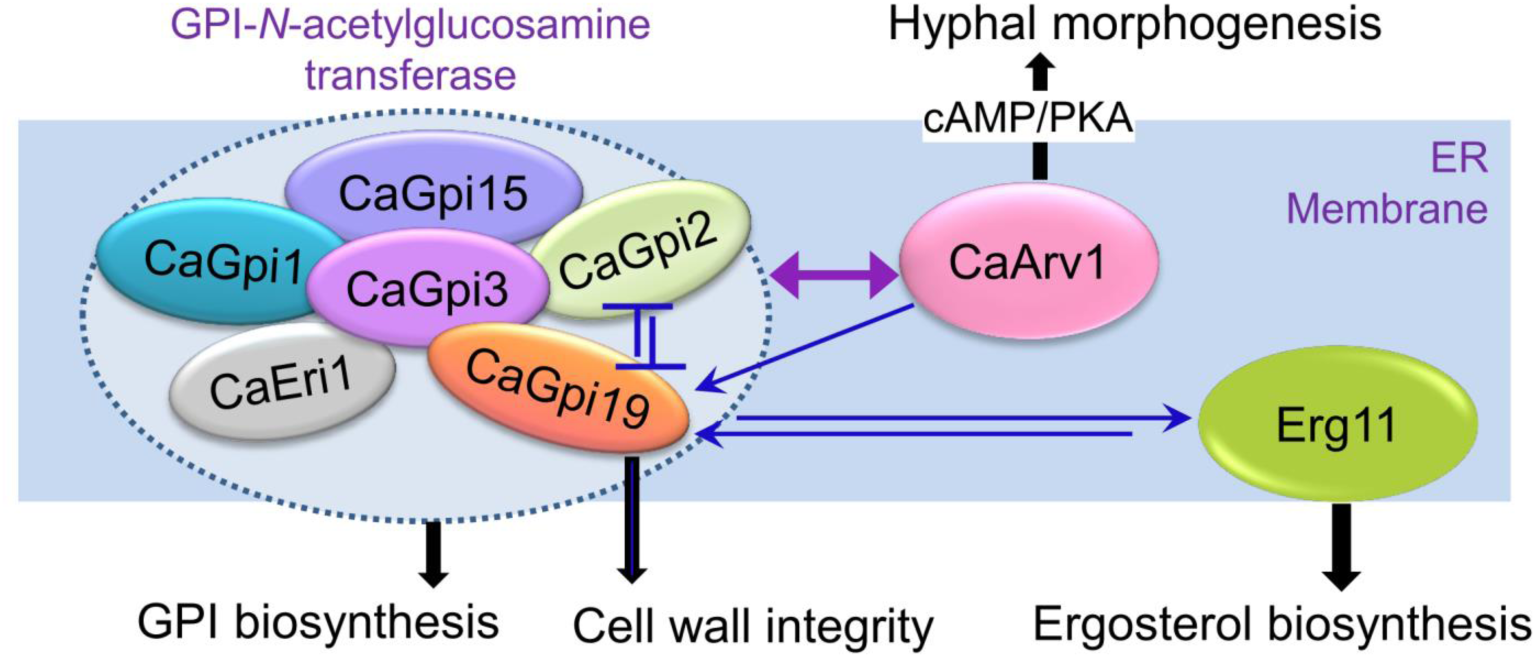
A model for the interaction of CaArv1 with GPI-GnT and its effect on the cell. CaArv1 physically interacts with the GPI-GnT and at least three of the GPI-GnT subunits (CaGpi2, CaGpi19 and CaGpi3) are in sufficient proximity to establish a physical interaction with it. CaArv1 also controls GPI-GnT activity via transcriptional regulation of the different GPI-GnT subunits and produces a cold-sensitive phenotype. In addition, the null strain of *CaARV1* shows cell wall integrity defects, and reduced ergosterol biosynthesis due to downregulation of Erg11. All of these phenotypes are reversed by overexpression of *CaGPI19* in this strain, suggesting that these phenotypes are all controlled via the GPI-GnT acting downstream of CaArv1. That a co-regulation exists between *CaGPI19* and *ERG11* has been previously reported [15]. Although *C. albicans* GPI-GnT can regulate the cAMP-PKA dependent hyphal morphogenesis pathway [18, 21], the hypofilamentous phenotype of *CaARV1* null strain is independent of GPI-GnT despite occurring in media that activate cAMP-PKA dependent filamentation. The double headed purple arrow represents physical interaction, the single-headed dark blue arrows represent transcriptional activation, and the dark blue flat headed arrows represent transcriptional repression.

Results from the co-localization, co-IP and AP-FRET experiments presented here also suggest a physical interaction between CaArv1 and the GPI-GnT (Figure 6), with at least three of the GPI-GnT subunits, CaGpi2, CaGpi3, and CaGpi19, being potential interacting partners. This interaction too could cause ‘activation’ of the GPI-GnT. The presence of CaArv1 is not crucial for the GPI-GnT activity *per se* since the null strain of *CaARV1* still possesses GPI-GnT activity and expresses GPI anchored proteins on its cell surface, although at levels lower than in the wild type cells. The reduction in GPI-GnT activity in the *CaARV1* deficient strains cannot be simply compensated by supplementation with ergosterol. Indeed, even GPI biosynthetic strains with upregulated ergosterol levels show reduced GPI-GnT activity [20]. The fact that overexpression of *CaGPI19* in the *CaARV1* null strain restores GPI-GnT activity to wild type levels even in the absence of CaArv1, suggests that an overall reduction in the levels of functional GPI-GnT available could be the major reason for the reduction in GPI-GnT activity seen in the *CaARV1* null strain rather than a loss of physical ‘activation’.

What then might be the reason for, or consequence of, the physical interaction between CaArv1 and the GPI-GnT in *C. albicans*? While this too needs further investigation, one possible hypothesis would be that the physical interaction allows CaArv1 and the first step of the GPI biosynthesis to rapidly and mutually respond to alterations in the sterol/ GPI biosynthesis, thus permitting *C. albicans* to efficiently adapt to external cues and environmental challenges.

We also observe that the null strain of *CaARV1* is cold-sensitive. A similar phenotype is reported in the *S. cerevisiae arv1Δ* cells along with accumulation of GlcN(acyl)-PI and a deficiency in mature GPI-AP production, which led the authors to suggest that membrane fluidity and, hence, flipping of the GPI intermediate from the cytoplasmic side to the ER lumen was affected in *arv1Δ* cells [2,3]. Overexpression of *GPI1, GPI2, GPI3* or *ERI1* can rescue the cold sensitive phenotype of *arv1Δ* cells [2]. As is seen from this study, the cold sensitive growth of *CaARV1* null, along with the GPI-GnT activity, is rescued by the overexpression of *CaGPI19*. Whether CaArv1 functions as a flippase and causes accumulation of GlcN(acyl)-PI remains to be seen. Our immunostaining studies for Als5, a GPI-AP on the fungal cell surface, clearly shows a reduction in the levels of GPI-APs present at the cell surface of a null strain of *CaARV1*. Since GPI-APs are required for virulence in *C. albicans*, it is not surprising that strains lacking functional CaArv1 are avirulent [5,8,23].

In summary, the results presented here expand the repertoire of roles that Arv1 performs in the cell. While its role in sterol and lipid homeostasis has been well recognized, its role in GPI biosynthesis and regulation are only just beginning to be documented in different eukaryotes. *C. albicans* Arv1 plays a significant role in regulating GPI biosynthesis by exerting control on the very first biosynthetic step. In doing so, it also regulates the drug response of the cell via this pathway. Simultaneously, and independently, it downregulates the cAMP-PKA pathway and hence controls the expression of hyphae-specific virulence factors, many of which are themselves GPI-anchored. An interesting corollary of these results is also that the first step of GPI biosynthesis is emerging in its own right as a major hub for multiple cross-talk events and regulations in *C. albicans*. On the one hand it interacts with and controls filamentation of *C. albicans* and on the other it interacts with sterol biosynthesis and regulates its drug response [19]. In addition, it now appears to independently interact with CaArv1, thus controlling as well as responding to lipid/ sterol homeostasis in the cell.

## METHODS

### Materials

The growth media components were purchased from Hi-Media (India), amino acids, dextrose, and other chemicals from Sisco Research Laboratories (SRL), glass beads from Unigenetics Instruments Pvt. Ltd., calcofluor white from Merck-Sigma-Aldrich (USA), NP-40 alternative, tunicamycin, HPTLC plates, ECL reagent (WBLUF0100), and PVDF membrane from Merck-Millipore (Germany), Protein A-Sepharose beads from Merck-Cytiva (USA), yeast transformation kit from G-Biosciences (USA), ER tracker^TM^ Green (BODIPY^TM^ FL Glibenclamide) from Invitrogen (Thermo-Fisher Scientific, USA), DNA gel extraction kit from Macherey-Nagel (Germany), trizol reagent, SYBR green and PCR master mix from Kapa Biosystems (USA), Dulbecco’s Modified Eagle Medium (DMEM) from Gibco (USA), PCR reagents from Genei (India) and restriction enzymes from either New England Biolabs (USA) or Fermentas (USA). UDP-[6-^3^H]GlcNAc was purchased from American Radiolabeled Chemicals Inc. (USA). Antibodies for Als5 [30], Gpi8 [31] and Gpi3 (for this study) were custom generated by Genei (India). Other antibodies were purchased either from Cell Signalling Technology (USA), GeneTex (USA), Thermo-Fisher Scientific (USA) or Genei (India). Primers were custom synthesized by GCC Biotech (India) or Eurofins Genomics (France).

### Strains and growth conditions

All the strains used in this study are generated in *C. albicans* BWP17 and are listed in Table 1. All the strains containing uridine (Ura) were grown either in YPD or synthetic defined (SD) medium lacking Ura (Ura^-^).

### Generation of heterozygous, homozygous null and reintegrant strains of *CaARV1*

All the primers used in this study are mentioned in Table S1. The *HIS1* selection marker was used to generate the heterozygous mutant (*Caarv1Δ*) by replacing one allele of *CaARV1* in the BWP17 strain of *C. albicans* while the null mutant (*Caarv1Δ/Δ*) was generated by replacing the surviving *CaARV1* allele with *ARG4* selection marker in the *Caarv1Δ* background. The mutants were confirmed by gene-flanking forward primer (FP) and reverse primer (RP), respectively, using the genomic DNA of the transformant colonies as template. For generating *Caarv1Δ/Δ-*P*_ACT1_-CaARV1,* the *CaARV1* gene was cloned into p*ACT1-GFP* vector [32] and then used to transform the *Caarv1Δ/Δ* strain. The transformants were confirmed by PCR with gene specific forward primer and *RPS1* locus reverse primer. Since this vector carries a *URA3* marker, the empty p*ACT1-GFP* vector alone was used to transform BWP17 and *Caarv1Δ* strains to obtain strains containing *URA3* at the *RPS1* locus for our studies.

### PCR based amplification of *CaARV1* gene

The forward and reverse primers (Table S1) designed using the sequence retrieved from CGD [33] were used to amplify the *CaARV1* gene.

### Growth assays

Primary cultures were grown on SD Ura^-^ medium for 16 h, 220 rpm at 30 °C. Secondary cultures were inoculated with cells corresponding to O.D._600 nm_ ∼ 0.2 in 60 ml SD Ura^-^ medium and incubated at 30 °C, 220 rpm. Similarly secondary culture with ketoconazole (0.15 µg/ml) was analyzed to examine drug sensitivity. O.D._600nm_ was measured every 2 h up to 24 h. Growth curves were plotted as O.D._600 nm_ versus time. Doubling times were calculated using the GraphPad Prism software.

### Spot assays

Secondary cultures were grown till mid-log phase using a 2% inoculum from a primary culture. Cells corresponding to O.D._600nm_ ∼ 0.1 were taken, and five-fold serial dilutions were prepared in 0.9% saline. Suspensions of 5 μl from each dilution were spotted on SD Ura^-^ agar plates with or without drugs, incubated at 30 °C. Images were taken using an iBright^TM^ CL1500 imaging system after every 12 h until the growth reached saturation.

### Sterol analysis by GC-MS

Estimation of sterol extraction and analysis were done as described in [15]. Briefly, cells from a secondary culture corresponding to O.D._600nm_ of ∼ 50 were lysed using acid-washed beads, the supernatant transferred to glass tubes and mixed with 9 ml of CHCl_3_:CH_3_OH (2:1 v/v). After incubating the sample for 2 h at 37 °C, the supernatant was discarded and 0.9% saline was added to the sample, followed by centrifugation. The bottom layer was taken into fresh pre-weighed tubes and incubated at 50 °C for 45 min. Once dry, the tubes were weighed again. To the sample 500 μl of CHCl_3_ was added and vortexed (2 x 1 min). Thereafter, 500 μl of 0.3 M methanolic KOH was added followed by vortexing. The sample was incubated at 37 °C for 1 h, then neutralized with 325 μl of 1M HCl, and further centrifuged at 300 rpm for 10 min. The lower layer was taken and dried in glass vials under a stream of N_2_ gas. For sialylation, BSTFA + TMCS (100 μl; incubation at 85 °C, 100 min) was used. Cholesterol externally added before the sterol extraction was used as the control for estimating extraction efficiency and normalization.

### Post-mitochondrial fraction (PMF) and GPI-GnT assay

PMF preparation and GPI-GnT assays were carried out as per previously established protocols [20]. Briefly, cells harvested from 250 ml secondary cultures of the *C. albicans* strains were lysed with acid-washed beads in 50 mM Tris-HCl buffer (pH 7.2) containing 10 mM EDTA. The lysate was centrifuged twice at 4 °C, first at 1000x g to remove the cell debris and then at 12000x g to obtain the PMF (supernatant).

PMF equivalent to 1500 µg total protein was incubated with UDP-[6-^3^H]GlcNAc (1 µCi) and tunicamycin (0.25 mg/ml) for 2 h at 30 °C. Heat-killed PMF was utilized as a negative control. The glycolipids were extracted in a solvent mixture of 10:10:3 chloroform/methanol/water, and dried under N_2_ gas (Grade-I). They were then partitioned into the upper butanol phase using water-saturated n-butanol:water (2:1). The butanol layer was aspirated out and dried again to remove the solvent. The glycolipids were resuspended in 10 µl of water-saturated n-butanol for spotting and resolving on HPTLC plates using 65:25:4 chloroform/methanol/water. The radiolabeled glycolipids were detected using a Bioscan AR-2000 TLC scanner and quantified with the WinScan software.

### Quantification of Als5 levels

Cells from a secondary culture at O.D._600nm_ ∼ 1.0 were harvested and subjected to incubation with anti-Als5 antibody (1:100) overnight at 4 °C on a rocker [30]. The cells were then washed thrice with 1X PBS buffer, and incubated for 2 h at room temperature in the dark with TRITC-labelled anti-rabbit secondary antibody (1:100). They were washed again with 1X PBS buffer, and resuspended in 50% glycerol. A 10 µl suspension was taken on glass slides and observed under a Nikon A1R HD25 confocal microscope. Data analysis was carried out using the Nikon NIS element 4000 AR analysis software.

### Filamentation assay

In order to induce hyphal growth in Spider medium, a 2% inoculum from a primary culture was used. Hyphal growth was observed after incubation at 37 °C for 90 min. To stimulate CO_2_-dependent filamentation, 4 ml of DMEM containing 40 mM sodium bicarbonate was inoculated with a 2.5% inoculum from the primary culture and incubated at 37 °C in the presence of 5% CO_2_ for 4 h. The cells were visualized under Nikon Eclipse TiE microscope. Manual counting of the various morphological forms including yeast, germ tubes, pseudohyphae, and hyphae was conducted, and the length of the hyphae was estimated using the Nikon NIS element 4000 AR analysis software.

### Calcofluor white (CFW) staining

A 2% inoculum from the primary culture was used to initiate a secondary culture which was grown for 6 h at 30 °C. Cells corresponding to O.D._600 nm_ ∼1.0 were stained with 100 µg/ml of CFW by incubation for 20 min on rocker shaker in the dark. The cells were pelleted down, washed thrice with 1X PBS, and resuspended in 50 µl of 80% glycerol. A 10 µl cell suspension was spotted onto a microscopic glass slide and visualized under a Nikon A1R HD25 confocal microscope. The data were analyzed using the Nikon NIS element 4000 AR software.

### Minimum inhibitory concentration for 80% growth inhibition (MIC_80_)

MIC80 assays were performed as reported previously [20]. Secondary cultures of the strains were grown for 6 h at 30 °C from 2% inoculum of their respective primary cultures. Equal aliquots of cell suspension (10^3^ cells/ 100 μl) were then plated in wells of a 96-well flat bottom microtiter plate. Two-fold serial dilutions of ketoconazole were added to all the wells and the plate incubated at 30 °C with mild shaking. After 24 h, O.D._600 nm_ was measured using a multi-plate reader (Thermo-Fischer Scientific, USA). The MIC_80_ was calculated as the concentration of drug where 80% inhibition of cell growth was seen.

### Transcript levels analysis using qPCR

RNA isolation and cDNA synthesis procedures were carried out on mid-log phase cells by a protocol previously reported [20]. Subsequently, qPCR analysis was conducted employing SYBR green PCR master mix along with gene-specific RT primers. The quantification of gene expression at the mRNA level was performed utilizing the comparative Ct method. The housekeeping gene *GAPDH* was employed as the internal control.

### Co-immunoprecipitation (co-IP)

Secondary culture of BWP17-*CaGPI2-6X-HIS*/*CaARV1-V5* strain was cultured till O.D._600 nm_ reached between 1 to 2. Cells were lysed in lysis buffer (10 mM HEPES-Na pH 7.5, 200 mM (NH_4_)_2_SO_4_, 5 mM MgSO_4_, 5 mM EDTA, 0.1% NP-40 substitute, 200 mM NaCl, 0.5% Triton-X, 10% glycerol). The cell lysate was centrifuged at 6000 rpm, 4°C for 10 mins and the obtained supernatant was incubated overnight with anti-V5 (#13202, Cell Signaling Technology) and Protein A-Sepharose beads at 4°C. The unbound supernatant was removed by centrifuging at 2000 rpm, 4°C for 1 min and beads were washed with lysis buffer three times. Bound proteins were eluted in SDS-PAGE sample buffer and samples for Western blotting was prepared by boiling at 95°C for 10 mins.

### Pull-down assays

Secondary cultures of BWP17-*CaGPI2-6X-HIS*/*CaARV1-V5* and BWP17-*CaGPI19-6X-HIS*/Ca*ARV1-V5* were grown in SD Ura^-^ until O.D._600 nm_ was between 1.0 to 2.0. The cells were harvested, lysed in lysis buffer (10 mM HEPES-Na pH 7.5, 0.1% NP-40 alternative, 200 mM NaCl, 0.5% Triton-X, 10% glycerol) using glass beads (10 x 1 min on vortex mixer, 1 min on ice). After centrifugation at 6000 rpm, 4 °C for 10 min, the supernatant was incubated with Ni^2+^-NTA beads at 4 °C overnight. The unbound supernatant was removed after centrifugation, the beads washed thrice with lysis buffer and the bound proteins eluted in SDS-PAGE sample buffer for analysis on 12% SDS-PAGE gels.

### Western blotting

The proteins were resolved on 12% SDS-PAGE gels, transferred onto PVDF membranes, blocked with 5% skimmed milk in PBS containing 0.05% Tween 20, and incubated overnight at 4°C with the primary antibody, either with anti-His (#SC-8036; Santa Cruz Biotechnology) at 1:500 dilution, anti-V5 (#13202, Cell Signaling Technology) at 1:1000 dilution, anti-CaGpi3 at 1:1000 dilution, or anti-β-actin (#GTX109639, GeneTex) at 1:1000 dilution. After extensive washing, HRP-conjugated secondary antibody (#114068001A, Genei) at a dilution of 1:2500 or 1:5000 was added, and the membranes incubated at room temperature for 2 h. The blots were developed on X-ray films using an ECL reagent.

### Super-resolution radial fluctuations (SRRF) Microscopy

Super-resolution of co-localization images from the Nikon A1R HD25 confocal microscope was achieved by using the NanoJ-SRRF plugin in ImageJ software as described previously [18]. Six radial axes within a 0.5 pixel ring radius were used with a radiality magnification of 5 for the analysis. SRRF analysis was carried out separately for each fluorophore by splitting the data obtained from the two channels (488 nm for FITC; 561 nm for TRITC). A temporal radiality average of 109 frames were used to obtain the temporal parameter. False color in ImageJ software was used to generate the final images.

### Acceptor-photobleaching Förster resonance energy transfer (AP-FRET)

CaGpi3 was detected by primary anti-CaGpi3 antibodies while His_6_ and V5 tagged proteins were detected by primary anti-His_6_ and anti-V5 antibodies in *BWP17-CaGPI2-6X-HIS/CaARV1-V5* and *BWP17-CaGPI19-6X-HIS/CaARV1-V5* respectively. Appropriate secondary antibodies were used (Goat anti-mouse IgG-FITC and Goat anti-rabbit IgG-TRITC) to generate the fluorescent FITC-TRITC donor-acceptor pairs. *CaGPI2-6X-HIS/CaRAS1* and *CaGPI8/CaRAS1* strains were used as the positive and negative control strains, respectively. AP-FRET was performed using a Nikon A1R HD25 confocal microscope using a previously standardized protocol [18]. FITC was excited with a 488 nm HeNe laser and TRITC with a 543 nm HeNe laser. Emission from FITC was collected using a 500-530-nm band-pass filter and that from TRITC was excited using a 550-590 nm band-pass filter. Cells labelled only with FITC-conjugated antibodies gave no bleed-through in the TRITC-channel. Excitation by the 488 nm laser produced no non-specific excitation in cells labelled with TRITC-conjugated antibodies alone. The TRITC label was photobleached by repeated irradiation with a 543 nm HeNe laser. Fluorescence emission from FITC was monitored before (pre-bleach) and after (post-bleach) photobleaching of TRITC in 20 × 20 pixels^2^ region of interests (ROIs) where the pre- and post-bleach image of the donor (Arv1-V5-FITC) aligned with that of the acceptor (for eg. CaGpi2-His_6_-TRITC). This was done by tracking their X-Y coordinates in ImageJ software. A minimum of 4 ROIs per cell was measured. The experiment was repeated at least three times for 4 cells of each strain and FRET efficiency (E) and donor-acceptor distance (*r*) measurements were calculated using the following two equations:

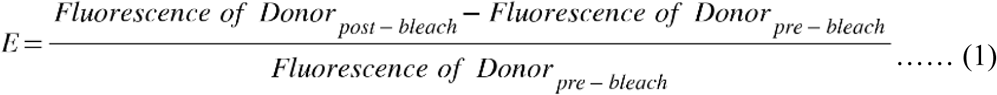

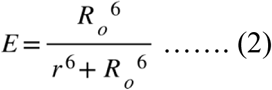

where, *R_o_*is the Förster distance (taken as 5.6 nm) for FITC-TRITC pair.

### Statistical analysis

All the analyses were performed on GraphPad Prism 10.0 software. unpaired T-tests were performed with Welch’s correction to calculate the statistical significance for pairwise samples. Two-way ANOVA with Tukey’s multiple comparison analysis was performed for multiple comparisons. Symbols */^#^ denote P values. The statistical significance are as follows: */^#^ ≤ 0.05; **/^##^ ≤ 0.01; ***/^###^ ≤ 0.001

## Abbreviations

AP-FRET: acceptor photobleaching-fluorescence resonance energy transfer
*ARV1*: *ARE2* required for viability1; GPI-glycosylphosphatidylinositol
cAMP-PKA: cyclic adenosine monophosphate-protein kinase A
CFW: calcofluor white
co-IP: co-immunoprecipitation
ER: endoplasmic reticulum
GC-MS: gas chromatography-mass spectrometry
GPI-AP(s): GPI-anchored protein(s)
GPI-GnT: GPI-N-acetylglucosamine transferase
GPIT: GPI transamidase
PI: phosphatidylinositol
PM: plasma membrane
PMF: post-mitochondrial fraction
SRRF: super resolution radial fluctuation

## Conflict of Interest

The authors have no conflict of interest to report.

## Author Contributions

MB, HS, YK, NT, UY, SS, SSaun, and AA performed the experiments. SSaun and AA participated as short-term project trainees on this project. MB and HS have contributed equally as first authors and YK and NT as second authors to this work. SSK conceptualized the project and brought in the funds. Data analysis/ interpretation and manuscript writing was by SSK, MB, HS and NT with specific inputs from the others.

## Acknowledgements

Parts of this work were funded by grants to S.S.K. from Department of Biotechnology (DBT) India (BT/PR29186/BRB/0/1726/2018) and Science and Engineering Research Board (SERB), India (CRG/2020/001649). SSK is also a recipient of the SERB-POWER Fellowship (SPF/2021/000097). M.B. was supported by Senior and Junior Research fellowships (SRF/ JRF) from University Grants Commission, India. Y.K. received a SRF, and U.Y. received Research Associate fellowship (45/10/2022-BIO/BMS), respectively, from the Indian Council of Medical Research. N.T. received a JRF from the Council of Scientific Research (CSIR), India. A.A. received a stipend from SSK’s SERB project and H.S. received a salary from the SERB/ SERB-POWER Fellowship grant of S.S.K. Confocal microscopy was performed at the Central Instrumentation Facility (CIF), School of Life Sciences (SLS), JNU. Facilities and research at SLS have been supported by grants from UGC-CAS, UGC-RNW, DBT-BUILDER, and DST-FIST.

## Notes

### Competing Interest Statement

The authors have declared no competing interest.

