## Supporting information for "Arv1 interacts with and regulates the first step of GPI biosynthesis in *Candida albicans*"

**Running Title: *C. albicans* Arv1 regulates GPI biosynthesis**

**Supporting Information:**

**Table S1: List of primers used in this study**

| **Primer name** | **Primer sequence** |
| --- | --- |
| FPCaARV1-HIS1 | 5'-ATGATCTGTATAGAATGTGGATATTCTAATATTGATTGTTTATATTCC AAGTATAAAAGCGGGGATCCTGGAGGATGAG-3' |
| RPCaARV1-HIS1 | 5'-ATTACAATGAAATGTATCAATTATATTCACAGATTCAAACCTATAT ACGTTGGATTTAACGGAATATTATGAGAAACT-3' |
| FPCaARV1-ARG4 | 5'-GAAGAGCTTACACAATCACTACCTATCATCATCATCACCATCCATT  TCATGT GGAATTGTGAGCGGAAG-3' |
| RPCaARV1-ARG4 | 5'-TCAGGTACAACTGTACGGAAACCACCTCAATATTTCAGTTCAGAATG  GGACTATTTCCCAGTCACGACGTT-3' |
| InFPCaGPI19 | 5'-GAAGTTTTAACATTACCTTTAAACGATATTCAT-3' |
| InFPCaARV1 | 5'-GAAGTTTTAACATTACCTTTAAACGATATTCAT-3' |
| LSFPCaARV1 | 5'-CGCAAGCTTATGATCTGTATAGAATGTGG-3' |
| LSRPCaRPS1 | 5'-AATAGAGAGAAACTATATTATACAC-3' |
| FPCaARV1HindIII | 5'-CGCAAGCTTATGATCTGTATAGAATGTGG-3' |
| FPGFP | 5'-CTAGCTTATTTGTACAATTC-3' |
| FPCaGPI19-6X- HIS-URA3 | 5'-AGTGGTGTTTGGGATTTGCCTATTACATTAGTGAATGATGTTCTATA TGAACATCACCACCACCATCACTAATAATAGGAATTGATTTGGATGG |
| RPCaGPI19-6X- HIS-URA3 | 5'-GGTATATATATACATATATAGTTTTTTGCTTTTCATTAATATTTGTGCA TTTTATTCATTATCATCATTATATAATTGGCCAGTCTTTTTC-3' |
| FPCaARV1-V5-ARG4 | 5'-ATTATATTCACAGATTCAAACCTATATACGTTGGATTGGTAGCCTATC CCTAACCCTCTCCTCGGTCTCGATTCTACGTAATACAATCATTT-3' |
| RPCaARV1-V5 | 5'-AGTTAATGATTTAGACGGGCCAATGATTGCATTGGATGGTGAAGAT GTATTGATAGTTAATCGAGATTTAACTTAAAATTGATTTTAAATTTTC-3' |
| RT FPCaGAPDH | 5'-CAGCTATCAAGAAAGCTTCTG-3' |
| RT RPCaGAPDH | 5'-GATGAGTAGCTTGAACCCAA-3' |
| RT FPCaERG11 | 5'-GTGGTGGTAGACATAGATGT-3' |
| RT RPCaERG11 | 5'-CCATCAATAGTCCATCTTAAA-3' |
| RT FPCaGPI1 | 5'-TCAGGGGATGACAATATTAAA-3' |
| RT RPCaGPI1 | 5'-CACGGAAATTTTTTGCC-3' |
| RT FPCaGPI2 | 5'-GGCCAATAGCATTTCTAAC-3' |
| RT RPCaGPI2 | 5'-CCATCACAAAAACAGACAAA-3' |
| RT FP-CaGPI3 | 5'-TGCTGAACCGGAAGAAAACT-3' |
| RT RP-CaGPI3 | 5'-TACATCTTTGCAACCGCATC-3' |
| RT FPCaGPI15 | 5'-CCACGATTATGGGCAGGTTA-3' |
| RT RPCaGPI15 | 5'-CGCGGTAAGAATTCTGGAAA-3' |
| RT FPCaGPI19 | 5'-CAAGAAGAAGAAGAAGGAGAA-3' |
| RT RPCaGPI19 | 5'-AAACACCACTTGGTGCCTTA-3' |
| RT FPCaERI1 | 5'-ACAGTCATCGCCAACATCCT-3' |
| RT RPCaERI1 | 5'-ATGGCCCACCACCAACATAC-3' |
| RT FPCaARV1 | 5'- ACAACGATGGTATCAGGG-3' |
| RT RPCaARV1 | 5'- TCGTGACAACTCGTAGTG-3' |

**Supporting Figures**

**Figure S1**

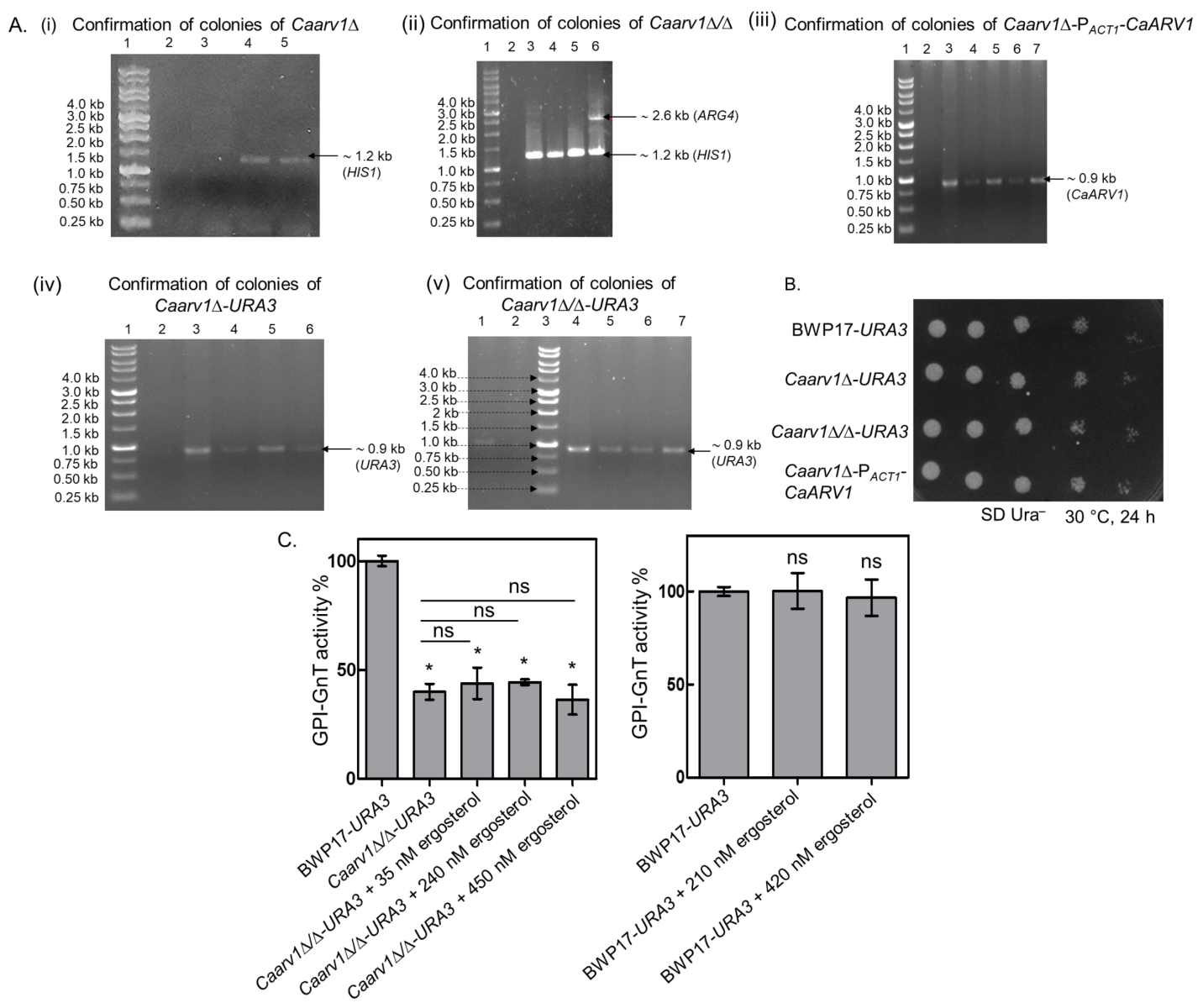

**Figure S1:** **Generation and confirmation of *CaARV1* mutant strains.** **A.** **Confirmation of *Caarv1∆, Caarv1∆/∆, Caarv1∆-*P*_ACT1_-CaARV1, Caarv1∆-URA3* and *Caarv1∆/∆-URA3*. (i)** The *HIS1* selection marker was used to generate the heterozygous mutant; it was confirmed by locus specific LSFPCaARV1 and RPCaARV1-HIS1 using the genomic DNA as template. Lane 1: 1 kb DNA ladder 2: negative control; Lane 3: negative colony; Lane 4,5: positive colonies of the *Caarv1∆* mutant. **(ii)** Homozygous null mutant was generated with *ARG4* marker in the *Caarv1∆* background. The mutants were confirmed by locus specific LSFPCaARV1 and RPCaARV1-ARG4 using the genomic DNA as template. Lane 1: 1 kb DNA ladder 2: negative control; Lane 3: genomic DNA of *Caarv1∆;* Lane 4,5: negative colonies; Lane 6: positive colony of *Caarv1∆/∆* mutant. **(iii)** Digested *pACT1-CaARV1* with StuI was used to generate *Caarv1∆-*P*_ACT1_-CaARV1* in *Caarv1∆* background*.* Strain was confirmed by primers such as FPCaARV1HindIII and LSRPCaRPS1. Lane 1: 1 kb DNA ladder, Lane 2: negative control; Lane 3: *pACT1-CaARV1,* positive control; Lane 4 & 6: negative colonies, Lane 5&7: positive colonies. **(iv)** StuI digested *pACT1-GFP* was used to generate *Caarv1∆-URA3.* Strain was confirmed by setting the PCR with the primers such as FPGFP and LSRPCaRPS1. Lane 1: 1 kb DNA ladder; Lane 2: negative control; Lane 3: *pACT1-GFP,* positive control; Lane 4-6: positive colonies. **(v)** StuI digested *pACT1-GFP* plasmid was used to transform the *Caarv1∆/∆* strain to generate *Caarv1∆/∆-URA3* strain. It was confirmed with PCR using FPGFP and LSRPCaRPS1 from genomic DNA of the transformants. Lane 1: *pACT1-GFP,* positive control; Lane 2: negative control; Lane 3: 1 kb DNA ladder; Lane 4-7: positive colonies*.* **B.** ***CaARV1* mutants show no growth defects on solid media**. *C. albicans* cells grown till mid-log phase were taken at O.D._600nm_ corresponding to 0.2 and serial dilutions were spotted on SD Ura^-^ solid media. Plates were incubated at 30 °C and images were taken at every 12 h till growth reaches saturation. **C. GPI-GnT activity is not restored in** ***Caarv1∆/∆-URA3* and in BWP17-*URA3* supplemented with ergosterol.** Post-mitochondrial fraction (PMF) of *Caarv1∆/∆-URA3* deficient in ergosterol was supplemented with specific amount of ergosterol to match the levels of WT BWP17-*URA3*; also increased amounts of ergosterol were used to carry out GPI-GnT activity. Different amounts of ergosterol were added to PMF of BWP17-*URA3* to carry out GPI-GnT activity. Averages and standard deviations are mean of three independent experimental sets. The statistical analysis was calculated using unpaired T-test with Welch’s correction.

**Figure S2**

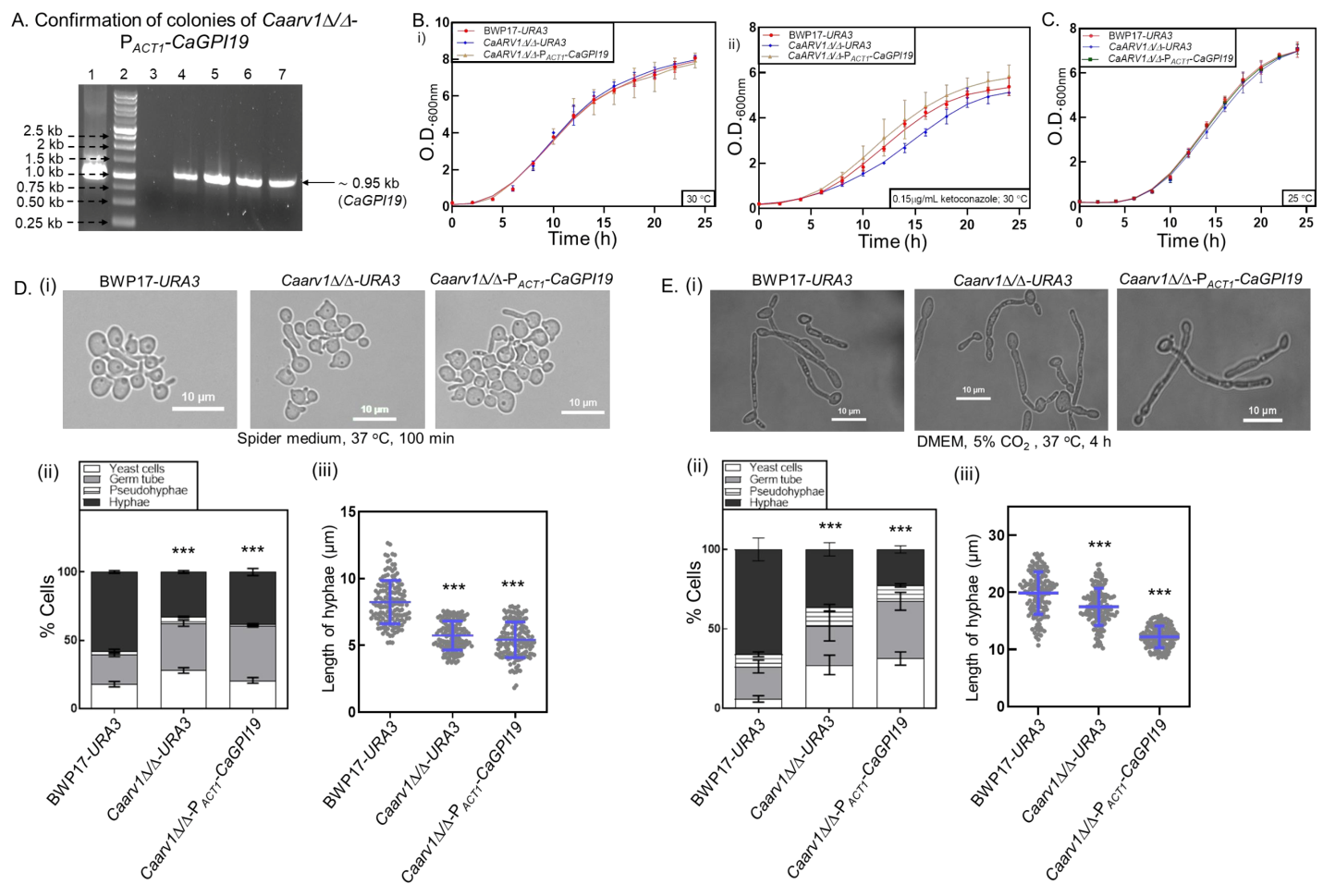

**Figure S2: Generation and confirmation of *Caarv1∆/∆-* P*_ACT1_-CaGPI19* strain and restoration in filamentation phenotype. A. Confirmation of *Caarv1∆/∆-*P*_ACT1_-CaGPI19.*** *pACT1-CaGPI19* was digested with StuI and used to generate *Caarv1∆/****∆****-*P*_ACT1_-CaGPI19* in *Caarv1∆/****∆*** background*.* Strain was confirmed by locus specific primers LSFPCaGPI19 and LSRPRPS1. Lane 1: *pACT1-CaGPI19,* positive control; Lane 2: 1 kb DNA ladder, Lane 3: negative control; Lane 4-7: positive colonies. **B. Azole sensitivity of *Caarv1∆/∆-URA3* is reversed upon *CaGPI19* overexpression.** Secondary cultures were grown at 30 °C in SD Ura^–^ liquid media using cells from a primary culture corresponding to O.D._600nm_ ~ 0.2. Growth was monitored every 2 h in the **(i)** absence and **(ii)** presence of 0.15 µg/ml ketoconazole. The doubling times obtained are given in Table 2. **C. *CaGPI19* overexpression reverses the cold sensitivity of *Caarv1∆/∆-URA3*.** Secondary cultures were grown at 25 °C in SD Ura^–^ liquid media using cells from a primary culture corresponding to O.D._600nm_ ~ 0.2. Growth was monitored after every 2 h. The doubling times obtained are given in Table 2. **D. Filamentation in Spider media reduces for *Caarv1∆/∆-* P*_ACT1_-CaGPI19* relative to *Caarv1∆/∆-URA3.* (i)** Representative images for each strain. **(ii)** Graph depicting number of yeast cells, germ tube, pseudohyphae and hyphae calculated manually for 100 cells each from 3 independent experiments. Averages and standard deviations are presented for hyphal cells only and was calculated using two-way ANOVA with Tukey’s multiple comparison test. **(iii)** Hyphal length quantification done for 50 cells each from 3 independent experiments. Averages and standard deviations are mean of three independent experimental sets. The statistical significance was calculated using the unpaired T-test with Welch’s correction. **E.** **Filamentation in DMEM + 5% CO_2_ reduces for *Caarv1∆/∆-* P*_ACT1_-CaGPI19* relative to *Caarv1∆/∆-URA3.*** **(i)** Representative images for each strain. **(ii)** Graph depicting number of yeast cells, germ tube, pseudohyphae and hyphae calculated manually for 100 cells each from 3 independent experiments. Averages and standard deviations are shown for hyphal cells only and was calculated using two-way ANOVA with Tukey’s multiple comparison test. **(iii)** Hyphal length quantification done for 50 cells each from 3 independent experiments. Averages and standard deviations are from three independent experimental sets. The statistical significance was calculated using the unpaired T-test with Welch’s correction.

**Figure S3**

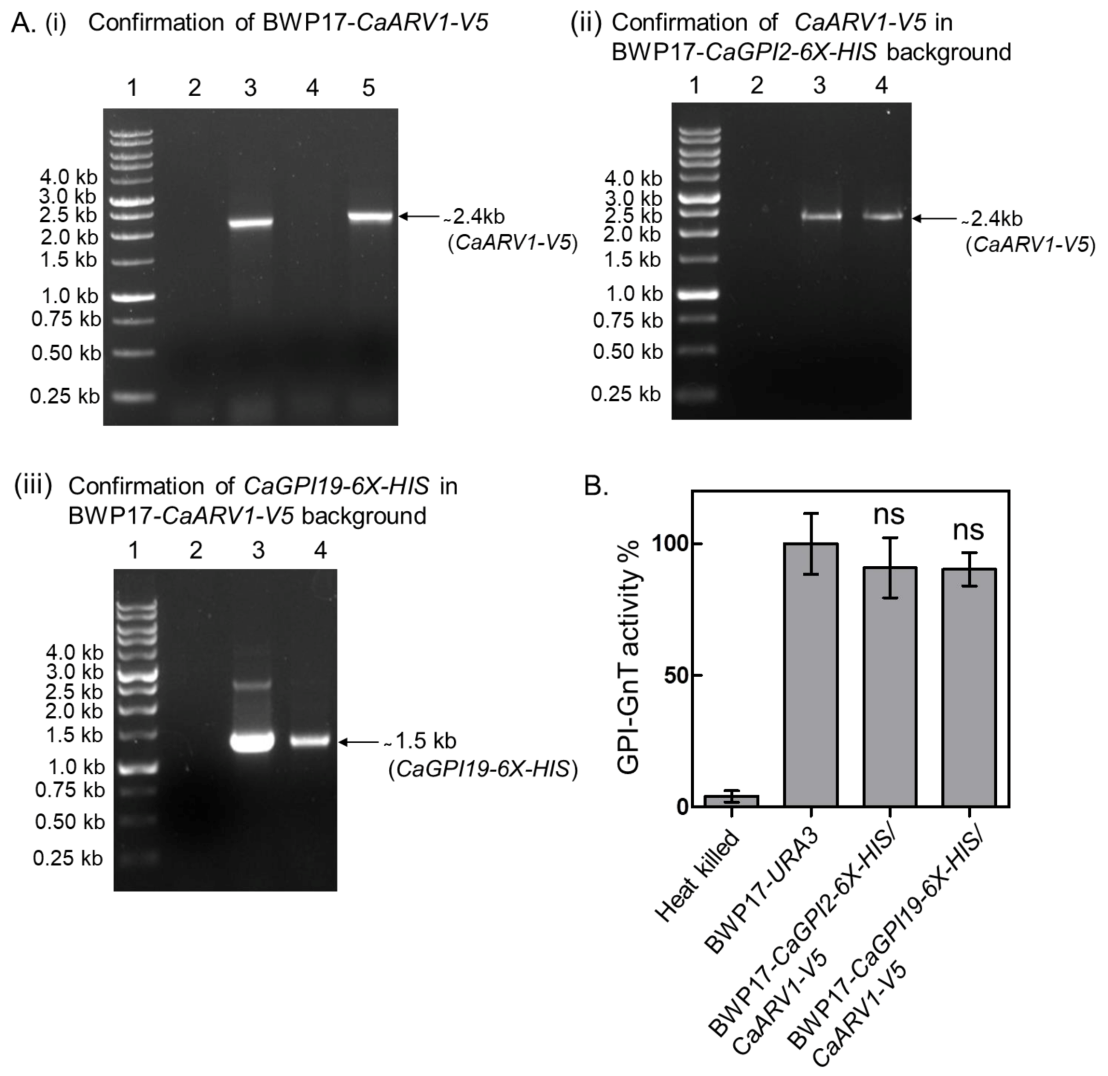

**Figure S3: Generation and confirmation of BWP17-*CaARV1*-*V5*****, BWP17*-CaGPI2-6X-HIS/CaARV1-V5* and BWP17*-CaGPI19-6X-HIS/CaARV1-V5* and the effect of the tag on GPI-GnT activity. A. Confirmation of BWP17-****Ca*ARV1*-*V5*, BWP17-*CaGPI2-6X-HIS/CaARV1-V5* and BWP17-*CaGPI19-6X-HIS/CaARV1-V5* strains*.* (i)** CaArv1 was tagged with V5-tag at its C-terminus by introducing the *V5* sequence in the primer used for amplification of *CaARV1*. *ARG4* was used as the selection marker. *CaARV1* was amplified using FPCaARV1-V5-ARG4 and RPCaARV1-ARG4 primers. The amplicon was used to transform BWP17-*URA3*. The colonies were confirmed by PCR using the primers InFPCaARV1 and RPCaARV1-ARG4. Lane1: 1 kb DNA ladder, Lane2: negative control, Lane 3&5: positive colonies, Lane 4: negative colony. **(ii)** Just as above**,** Arv1 was tagged with V5-tag at its C-terminus using ARG4 selection marker; it was amplified using FPCaARV1-V5-ARG4 and RPCaARV1-ARG4. The amplicon was used to transform BWP17-*CaGPI2-6X-HIS*; colonies obtained were confirmed by PCR using the primers InFPCaARV1 and RPCaARV1-ARG4 primers. Lane1: 1 kb DNA ladder, Lane2: negative control, Lane 3-4: positive colonies. **(iii)** Similarly, CaGpi19 was tagged with 6X-His tag at its C-terminus using *URA3* selection marker, it was amplified using FPCaGPI19-6X-HIS-URA3 and RPCaGPI19-URA3. The amplicon was used to transform BWP17-*CaARV1-V5*; colonies obtained were confirmed by PCR using the primers InFPCaGPI19 and RPCaGPI19-URA3. Lane1: 1 kb DNA ladder, Lane2: negative control, Lane 3-4: positive colonies. **B.** **Tagging of BWP17*-CaGPI2-6X-HIS/CaARV1-V5* and BWP17*-CaGPI19-6X-HIS/CaARV1-V5* does not affect GPI-GnT activity.** PMF of BWP17-*CaGPI2-6X-HIS/CaARV1-V5* and BWP17-*CaGPI19-6X-HIS/CaARV1-V5* strains were incubated with UDP-[6-^3^H]GlcNAc donor substrate and activity was compared with that of BWP17-*URA3.* Averages and standard deviations were calculated from three independent experiments. The statistical analysis was calculated using unpaired T-test with Welch’s correction.
